# Assessing reinforcement learning and vigor with an online decision-making battery

**DOI:** 10.64898/2026.10.30.763028

**Authors:** Štěpán Wenke, Tobias Granwald, Alessandro Ferrante, Martin Nilsson, Ingrid Ekström, Goran Papenberg, Marc Guitart-Masip

## Abstract

Reinforcement learning and vigor are two fundamental aspects of decision-making modulated by dopamine. Although vigor is highly relevant for mood disorders, it has been overlooked by previous batteries characterizing decision-making abilities. We developed a test battery with two RL tasks (RL/working memory, go/no-go task) and two vigor tasks (physical effort discounting, foraging task) and administered it to an online community sample (*N* = 162). We used computational modeling and factor analysis to identify two factors reflecting RL and vigor. The RL factor reflected participants’ general decision-making ability, while vigor corresponded to a motivational propensity to obtain rewards. Both factors demonstrated moderately strong test-retest reliability (ICC(2,1) = .65 and .67, respectively) over a one-to-two-week interval. Moreover, both factors were meaningfully related to self-report scales. The RL and vigor factors were negatively and positively associated with mania, respectively, although further analyses revealed that vigor was not fully independent of RL. Neither factor was significantly associated with depression symptoms. Collectively, we present preliminary evidence that vigor may represent a separate factor from RL in line with the putative dopamine hypothesis, paving the way for future longitudinal assessments of RL and vigor in mood disorders.

## Introduction

Decision-making is fundamental to our day-to-day life. We pick the appropriate clothes from our closet in the morning, choose which route to take to work, and what groceries to buy on the way home. Unsurprisingly, making advantageous decisions is therefore a prerequisite for well-being (Parker & Fischhoff, 2005), and decision-making processes are often altered in psychiatric disorders such as bipolar disorder (Murphy et al., 2001; Ramírez-Martín et al., 2020) or depression (Alexander et al., 2017; Scholl & Klein-Flügge, 2018; Treadway & Zald, 2011). Tasks assessing decision-making processes and learning offer a putative window into cognition by capturing ecologically relevant features of behavior. Assessing these abilities can therefore go beyond the traditional classification systems of mental disorders, which rely on clinical observation of clusters of symptoms rather than underlying pathophysiology (Hyman, 2007; Lee, 2013; Pizzagalli, 2014). Thus, the parameters obtained from these tasks may provide a way to mechanistically understand symptomatology in mood disorders (Huys et al., 2008; Montague et al., 2012; Scholl & Klein-Flügge, 2018).

Before introducing these tasks into psychiatric practice, it is of central importance that the psychometric properties of novel assessment tools are well established (Mkrtchian et al., 2023; Paulus et al., 2016). However, low reliability has been a prevalent issue in decision-making research (Brown et al., 2020; Pike et al., 2024; Schaaf et al., 2024; Zorowitz et al., 2025) but see also (Pike et al., 2026). One potential reason for this issue may be that the tasks used are not sensitive enough to reflect the ecological features of day-to-day processes (Scholl & Klein-Flügge, 2018). While past studies have demonstrated differences in key features of decision-making in patients using single tasks (Mukherjee et al., 2020; Rupprechter et al., 2018), it is challenging to draw any conclusions how these features are related to symptomatology due to the heterogeneity of tasks used (Pike et al., 2026). Decision-making is a multi-faceted construct composed of many underlying processes such as learning, valuation, or updating, collectively often referred to as reinforcement learning (RL) (Sutton & Barto, 2018). It is unlikely that capturing all these processes can be achieved using a single task (Scholl & Klein-Flügge, 2018). Moreover, it remains unclear which of these facets are most impacted by mood disorders (Mukherjee et al., 2020). Conducting studies with multiple tasks and forming composite scores, as opposed to estimating a mean performance from a single task, has great potential to increase the reliability and generalizability (Buelow et al., 2024; Endrass & Ullsperger, 2021; Granwald et al., 2025; Judd et al., 2024; Moutoussis et al., 2021; Parker & Fischhoff, 2005). A few studies have constructed task batteries exploring different dimensions of decision-making in the same participants (Mkrtchian et al., 2023; Moutoussis et al., 2021; Mukherjee et al., 2020; Parker & Fischhoff, 2005; Pooseh et al., 2017). One study with 7 decision-making tasks and 32 parameters found a single common factor reflecting a general decision-making ability, also termed decision acuity (Moutoussis et al., 2021). Decision acuity was stable over 18 months and was negatively associated with a general psychopathology factor, but it was unrelated to mood symptomatology. In another recent study by (Mukherjee et al., 2020), the authors constructed a comprehensive battery of 9 decision-making tasks and using factor analysis found that depressed individuals exhibited impaired learning, valuation, and patience. Further, the authors used machine learning to predict depressed status with 72% accuracy using data of task performance across all tasks, although no individual task predicted depression scores. While these studies demonstrate the importance of studying several tasks simultaneously to uncover dimensions of decision-making, they fail to show which, if any, of these aspects is related to mood psychopathology. Moreover, these studies leave out a crucial aspect, namely the vigor with which actions are executed.

Vigor is a crucial aspect but often left-out feature of decision-making (Niv, 2009). To optimize reward, animals and humans need to learn the value of available choices but also decide the vigor with which to execute those (McClure et al., 2003; Niv et al., 2007). This motivational aspect of decision-making has been suggested to reflect a dynamic cost-benefit analysis of expending energy to gain reward (Beierholm et al., 2013; Guitart-Masip et al., 2011; Niv et al., 2007; Wang et al., 2013) and is typically altered in neurological and psychiatric conditions (Cousins et al., 2009; Dunlop & Nemeroff, 2007; Treadway & Zald, 2014). For instance, lower vigor is associated with perceived lower reward in the environment and may be impaired in depression (Bonnelle et al., 2015; Bustamante et al., 2024, 2024; Huys et al., 2015; Treadway et al., 2009). As such, any account of decision-making that aims to predict psychopathology needs to also encompass vigor. To our knowledge, no previous study has included tasks measuring both RL and vigor. The current study aims to fill this gap by designing and testing a decision-making battery with tasks targeting both aspects.

Together, RL and vigor are two complementary aspects of decision-making which are likely altered in psychiatric disorders (Cousins et al., 2009; Dunlop & Nemeroff, 2007; Treadway & Zald, 2014) and putatively distinctly modulated by the dopaminergic system (Mohebi et al., 2019). On one hand, prior research points to the involvement of dopamine in RL via phasic dopamine signaling putatively representing the difference between the expectation and the outcome, termed reward prediction error (Collins & Frank, 2014; Montague et al., 1996; Schultz, 2016, 2019; Schultz et al., 1997). On the other hand, slower tonic dopamine has been associated with vigor (Beierholm et al., 2013; Berke, 2018; Collins & Frank, 2014; Hamid et al., 2016; Niv et al., 2007; Salamone & Correa, 2012). Specifically, it has been suggested that the baseline tonic dopamine levels reflect the average reward rate and regulate vigor by signaling an opportunity cost per unit time where time spent doing nothing results in potential foregone rewards (Niv et al., 2007). Moreover, ample evidence suggests that dopamine is involved in various psychiatric conditions, including depression (Dunlop & Nemeroff, 2007; Treadway & Zald, 2011) and mania (Cousins et al., 2009). Although the functional distinction between phasic and tonic dopamine may not be that straightforward (Hamid et al., 2016; Satoh et al., 2003), their possible selective effects on RL and vigor further justify the need to quantify both to understand decision-making and its relation to psychopathology.

In this preregistered study, we built and assessed a battery with two tasks measuring RL and two tasks measuring vigor that can be used to investigate the distinct contributions of these aspects of decision-making to psychopathology. We employed computational modelling to parametrize key cognitive mechanisms of interest in each task and used factor analysis to derive a composite score for RL and vigor based on a reduced selection of parameters. We also assessed the stability of the factors over a one-to-two-week interval. Furthermore, we investigated predictive validity of our newly developed online battery by assessing the association with mania and depression symptoms as well as apathy. Finally, we did an exploratory latent profile analysis to explore whether our data suggest distinct profiles of RL and vigor with different levels of mania, apathy, and depression symptoms.

## Methods

This study was preregistered before the start of the data collection at OSF: https://osf.io/z27nh/. The reported methods follow the preregistration and any deviations from it are clearly stated.

### Procedure

#### Ethical approval

According to the Swedish Act concerning the Ethical Review of Research Involving Humans (2003:460), ethical approval was not necessary for this study due to the nature of online data collection. The data were fully anonymized, containing no personal data or information that could be used to identify individuals. Moreover, the experiment did not pose any mental or physical burden that could affect the participants outside the study.

#### Study design

The study involved three sessions. Participants were asked to read and accept the consent form, read the general instructions, and complete the Go/No-Go Task. Upon completion, participants were compensated with a fixed bonus of £4, in addition to the performance bonus of up to £1.5. Session 1 and Session 2 were identical, separated by 5-14 days. Each of these sessions included 4 tasks and 6 questionnaires. Upon completion, participants were compensated with a fixed bonus of £14 and a performance bonus up to £6.

#### Data collection

Participants were recruited from the platform Prolific.com (Prolific). Out of a pool of 155,804 eligible participants (at the time of writing the preregistration), 306 were recruited to participate in Session 0. The experiment was hosted on a server http://pavlovia.org (Pavlovia; Open Science Tools, Nottingham, UK) and all tasks programmed in JavaScript with the help of a toolbox jsPsych version 7.3.3 (de Leeuw et al., 2023).

### Participants

#### Inclusion and exclusion criteria

The inclusion criteria were broad: age 18-60, fluency in English and no participation in either of our pilot studies. We did not apply any other exclusion criteria prior to Session 0. We invited participants from Session 0 to Session 1 only if they 1) performed above chance level (>= 55%) and 2) responded with “go” less or equal than in 90% of all responses. These criteria were used in a previous study using the same task (Zorowitz et al., 2025). To control for participants’ engagement online (Albert & Smilek, 2023), we embedded attention checks in each of the tasks and questionnaires. Participants were invited to a subsequent session only if they did not fail more than 2 attention checks in the same task or questionnaire. The target sample was 200 and 100 participants for Session 1 and 2, respectively. We chose this sample size based on the pilot studies in order to be able to detect correlations of at least .30, with α = .05 (two-sided) with power .80. For the details of the sample size rationale and demographic characteristics, please refer to Supplementary Materials.

After the data collection was completed, we inspected the data quality. If a participant did not fulfill any of the data quality criteria, they were excluded from the main analysis. The criteria were preregistered as follows: 1) exhibiting a below chance-level performance (<33% proportion of correct choice) in the RLWM Task, 2) exhibiting a below chance-level performance (<55% proportion of correct choice) and perseverance in “go” responses (>90% of all responses) in the GNG Task, 3) accepting every effort offer in the Effort task and 4) having the average exit threshold 3 standard deviations above the mean in the sample in the Foraging Task.

After applying these criteria for each Session, we obtained a final sample of 162 and 71 participants for Session 1 and Session 2, respectively. The schematic procedure of the study design, with inclusion and exclusion criteria, is shown in Figure 1.

**Figure 1.**
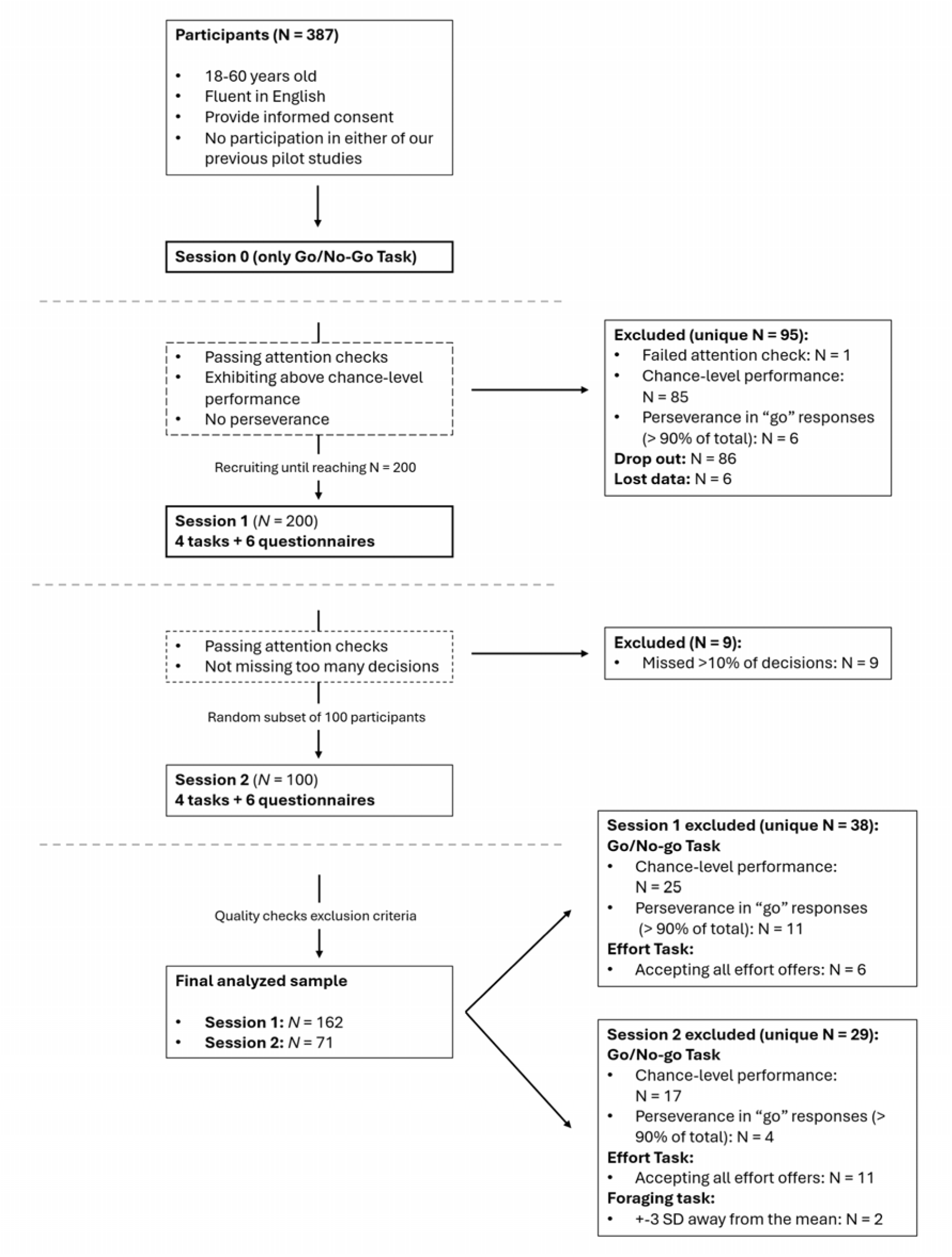
The schematic overview of the experimental procedure. *Note*. The experiment consisted of three sessions. The aim of the initial Session 0 was used to assess performance-based levels in the GNG Task and invite further eligible participants who passed the above-chance-level and non-perseverance criteria, as previously done for this task (Zorowitz et al., 2025).

#### Sample characteristics

Our sample from Session 1 included 162 participants (41.98% females, 57.41% males), while the sample at the re-test (Session 2) included 71 participants (40.85% females, 57.75% males).

#### Tasks

To capture RL, we employed a version of the reinforcement learning and working (RLWM) task (Collins & Frank, 2012; Master et al., 2020), and a valence Go/No-Go (GNG) task (Guitart-Masip et al., 2012; Zorowitz et al., 2025). These tasks measure value learning and updating independently from confounding cognitive mechanisms, namely working memory (Collins & Frank, 2012; Master et al., 2020) and affective bias (Guitart-Masip et al., 2012). For the vigor domain, we opted for a physical effort discounting Task (Bustamante et al., 2023; Lockwood et al., 2021) that quantifies willingness to exert physical effort to obtain variable rewards; and a foraging task (Harhen & Bornstein, 2023) that quantifies patch-leaving behavior: to exploit or explore. In the foraging task, participants had to decide when to leave a patch by estimating the reward distribution in the current patch, the overall richness of the environment, and the opportunity cost of time which is not spent harvesting (Addicott et al., 2017; Charnov, 1976; Lloyd et al., 2023; Stephens & Krebs, 1986). According to the Marginal Value Theorem (Charnov, 1976), an agent chooses to leave a patch when the expected reward of the current one falls below the average reward rate across all possible patches. As such, we hypothesized that patch-leaving thresholds (exit threshold) reflect the experienced average reward rate in the Foraging task, which together with vigor costs has been suggested regulate the level of vigor(Niv et al 2007). Supporting this interpretation, human participants exhibited higher exit thresholds after the administration of the dopaminergic precursor L-Dopa (Constantino et al., 2017) and dopamine agonist cabergoline (Le Heron et al., 2020). Similarly, higher dopamine synthesis capacity as measured with PET predicted higher exit thresholds (Ianni et al., 2023). Likewise, increased willingness of exerting effort to obtain reward is understood as a manifestation of vigor positively modulated by dopamine (Chong et al., 2015; Le Bouc et al., 2016; Le Heron et al., 2018; Salamone et al., 2007; Salamone & Correa, 2002, 2012). Taken together, we expected that the shared variance between effort discounting, reflecting inverse motivational propensity, and foraging exit threshold is a manifestation of vigor.

Here we provide a brief description of each task in the battery. Based on past studies, we implemented gamification elements in each task with the intention to increase engagement (Floryan et al., 2020; Hamari et al., 2014; Sailer et al., 2017; Verdejo-Garcia et al., 2021), performance (Wiley et al., 2024), and enjoyment (Friehs et al., 2020; Verdejo-Garcia et al., 2021; Wiley et al., 2024). For more detailed information about number of stimuli, trials and particular modifications we made to the adapted tasks, see the Supplementary Materials.

### Reinforcement Learning/Working Memory Task (RLWM Task)

This task was designed to disentangle slow RL mechanism from fast WM mechanisms during learning (Collins et al., 2014; Collins & Frank, 2012; Master et al., 2020). We programmed a version of the task based on the original study (Collins & Frank, 2012). Participants learned stimulus-response mappings for pictures from various categories, associating each with one of three response keys on the keyboard (*q, w,* or *e*). The stimuli provided no inherent cues about the correct response, and participants needed to learn the correct associations through trial and error. Feedback was deterministic (i.e., always accurate) and displayed after each response.

#### Go/No-go Task (GNG Task)

For the Go/No-go task we used openly available code (Zorowitz et al., 2025) and adapted it to our study. The task quantifies RL in the presence of a hard-wired motivational bias that interferes with RL when withholding a response (“No-go”) is required to obtain a reward (de Boer et al., 2019; Guitart-Masip et al., 2012, 2014; Swart et al., 2017). Participants were observing robots moving on a conveyor belt and, for each robot, learned whether to repair it (go response) or not (withhold response) based on probabilistic feedback. In the current study, we only used rewarded stimuli resulting only in go to win and no-go to win conditions. Based on pilot data, we observed that the parameters of the punishment conditions did not contribute to the RL factor and had worse parameter recovery indices. By removing the punishment conditions, we reduced the testing time while keeping the recovery of the retained parameter of interest high.

#### Effort Discounting Task (Effort Task)

The task quantifies the physical effort that one is willing to exert for increasing units of reward. For this study, we developed a new gamified task based on previous tasks quantifying physical effort discounting (Lockwood et al., 2021; Treadway et al., 2009). In our task, participants (shown as a goblin) competed for coins against a minotaur with the objective of getting as many coins as possible. In each trial, participants chose between taking a walk (rest option) to get a small reward or challenging a minotaur (effortful option) to obtain a bigger reward but requiring higher effort. Effort was operationalized as keyboard presses.

Due to programming error in our code, the training phase was shortened, and participants did not have the opportunity to experience all different levels of effort before the actual task started. Getting familiarized with the different levels of effort during training was initially planned and tested in our pilots.

#### Foraging Task

For the Foraging task, we used openly available code (Harhen & Bornstein, 2023) and adapted it to our study. In this task, participants mined gems from planets in space. In each trial, they could choose to either continue harvesting (digging for more gems) or leave (travel to another planet with replenished gems). The number of gems obtained decreased with each successive harvest on the same planet. The depletion rate was drawn from a beta distribution with a mean of 0.88 and variance of 0.01. Traveling to a new planet provided access to a replenished environment. However, it required time to get there (travel time). Therefore, participants needed to make a trade-off between extracting diminishing gems from an overmined planet and the opportunity cost of traveling to a new one.

At the end of the task, the total number of collected gems was converted into a performance bonus. According to the Marginal Value Theorem (MVT) (Charnov, 1976), participants should choose to leave a planet when the expected reward from the next dig falls below the average expected reward across all planets. We quantified participants’ average expected rewards by measuring the reward value when they exit the planet (exit threshold). In line with MVT, the exit threshold reflects the perceived average reward rate of the environment, which is suggested to be encoded by tonic dopamine levels (Niv et al., 2007), as supported by experimental data (Constantino et al., 2017; Ianni et al., 2023; Le Heron et al., 2020). The exit threshold was calculated as the average of the last two observed rewards before leaving the planet (Constantino & Daw, 2015; Lenow et al., 2017) and did not require the use of any computational modelling.

#### Questionnaires

We included three questionnaires in the main analysis based on preregistration. These questionnaires are introduced here while the description of the other questionnaires included in our battery can be found in Supplementary materials.

#### Apathy Motivation Index (AMI)

AMI is a brief questionnaire designed to quantify apathy in a healthy population (Y. S. Ang et al., 2017). Based on factor analysis, the authors found three different clusters of apathy and labeled them as behavioral activation apathy (BA), social motivation apathy (SM), and emotional sensitivity apathy (ES). The scale contains a total of 18 statements (each subscale contains 6 statements). Participants rated how true each statement was based on the preceding two weeks: ‘completely untrue’, ‘mostly untrue’, ‘neither true nor untrue’, ‘quite true’, and ‘completely true’. Each statement is negatively scored such that higher score represents greater apathy. Example statements are: “When I decide to do something, I am motivated to see it through to the end” from the BA subscale, “I suggest activities for me and my friends to do” from the SM subscale or “I feel awful if I say something insensitive” from the ES subscale. The scale exhibits good test-retest reliability (r = 0.83) and satisfactory internal consistency (α = .77).

#### The Altman Self-Rating Mania Scale (ASRM)

ASRM is a brief self-rating scale, compatible with DSM-IV criteria, to measure manic symptoms for research and clinical purposes (Altman et al., 1997). The scale contains 5 items concerning positive mood, self-confidence, sleep patterns, speech, and activity level. For each item, participants pick a statement that best describes how they have been feeling for the past week. The scale can distinguish patients experiencing mania from those not experiencing it. A cutoff score of 6 points or more indicates a manic patient (sensitivity of 85.50% and specificity of 87.30%). Furthermore, the scale has good test-retest reliability (r = .86, p < .001) and satisfactory internal consistency (α = .79).

#### Center for Epidemiologic Studies Depression Scale (CES-D)

CES-D is a short structured self-report scale designed to measure depressive symptoms in the general population (Radloff, 1977). It contains 20 statements and four possible answers to pick from based on the frequency during the past week (rarely or none of the time, some or a little of the time, occasionally or a moderate amount of time or most or all of the time). For example, there are statements such as “I felt sad.”, “I felt lonely” or “I thought my life had been a failure”. The scale was designed to measure the current depressive symptomatology, which is considered to vary over time and therefore only a moderate level of test-retest reliability was estimated (r = .55). However, the scale has a high internal consistency (α = .85) and excellent discriminant validity when correlating CES-D with other self-report scales (Radloff, 1977).

### Statistical and data processing

Computational modelling was carried out in MATLAB R2024a version 24.1. using built-in functions and custom scripts. All data processing and statistical analyses were conducted in R version 4.5.2 (R Core Team, 2025) using the following packages and libraries: *tidyverse* (Wickham et al., 2019), *dplyr* (Wickham et al., 2023) and *tidyr* (Wickham et al., 2025) for data manipulation and wrangling such as filtering, extracting and sorting data; *psych* (Revelle, 2015), *easystats* (Lüdecke et al., 2022), *ez* (Lawrence, 2026), *lme4* (Bates et al., 2015) and *emmeans* (Lenth & Piaskowski, 2025) for descriptive tables and inferential statistics such as ANOVA, regressions and post-hoc tests; *tidySEM* (Van Lissa, 2019) for carrying out LPA; *rempsyc* (Thériault, 2023) for creating APA-formatted tables; *broom* (Robinson et al., 2026) and *jsonlite* (Ooms, 2014) for data cleaning and *ggplot2* (Wickham, 2016) for plots and data visualization.

### Computational modeling

#### Models

Performance on each task except for the foraging task was modelled using standard RL or effort-discounting models previously utilized in literature or established as best models in pilot studies. Individual computational parameters were estimated using hierarchical Bayesian methods (Huys et al., 2011) (refer to Parameter estimation in the Supplementary Materials for more details). Below we describe which models were employed in each task.

For the details on parameter estimation, model comparison, parameter reliability, parameter distribution and other details, refer to Supplementary Materials for details of a specific model used in the respective task.

#### RLWM Task

The utilized model is composed of two mechanisms. **The first mechanism (RL)** is a Q-learning model (C. J. C. H. Watkins & Dayan, 1992) updating the learned Q-value for the selected action *a* given the stimulus *s* after receiving feedback on trial *t* regarding the reward outcome *r_t_* (1 for correct, 0 for incorrect):

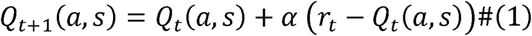

The free parameter α is the learning rate constrained between 0 and 1 that determines the extent to which the outcome influences the update of Q-values.

The RL mechanism generates choices probabilistically with a higher probability of choosing actions with higher Q-values based on a SoftMax choice function:

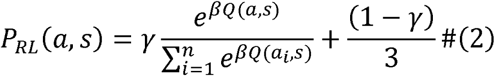

Here, β is the inverse temperature parameter controlling how deterministic choices are. The β parameter was fixed at 10 as previous research has shown that having both β and α as free parameters leads to a worse parameter identification (Master et al., 2020). The action set comprised of *n* = 3 responses, for the three possible actions *a_i_* participants could take (q, w, and e key on the keyboard). All Q-values were initialized at 1/n. To account for random choices and attentional lapses, we added a lapse rate estimated as the free parameter γ. It was constrained between 0 and 1.

**The second mechanism (WM)** is a fast but capacity-limited mechanism that stores the weights between the stimuli and actions, *W(a,s).* The weights are initialized at W_0_ = 1/n in the same way as Q-values in the RL module.

The weights are updated as follows:

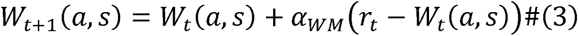

The where *α_WM_* = 1. Therefore, the action weights prescribed by the WM module capture a perfect retention of the previous trial’s stimulus value such that *W_t+_*_1_*(a,s)= r_t_* . The WM module allows for rapid learning and retrieval of stimulus–action associations with deterministic outcomes, unlike the slower, incremental learning in the RL module.

To model the information decay within the WM modules, we assumed forgetting which caused the WM weights to decay to their starting values at each trial according to:

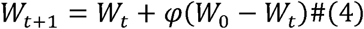

The decay parameter *φ* was constrained between 0 and 1.

As the RL module, the WM module utilized these weights to generate choices probabilistically with the SoftMax choice function with added lapse rate, using the same β and γ parameters as in the RL module.

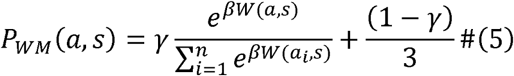

Choice between the RL and WM module was parametrized by the WM capacity parameter *K* and a WM reliance parameter ρ. The overall choice policy was defined as a mixture using WM weight *W* = ρ*min(1, K/ns):

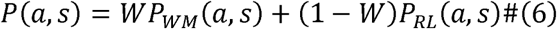

The free parameter ρ was constrained between 0 and 1 and captured the upper bound of participants’ tendency to use WM, as opposed to RL, when set size (ns) was within the capacity of WM. As set size increased over WM capacity, reliance decreased proportionally to the set size. The WM weight then contributed to the choice policy based on the capacity and the subject’s reliance on WM.

We also estimated a positive learning bias (*bias*) to capture the extent to which participants neglect negative outcomes. For negative prediction errors, when δ < 0, the learning rate α is reduced by α = (1 – bias)*α. The *bias* parameter was constrained between 0 and 1. Values near 1 indicate neglect of negative feedback, while values near 0 indicate unbiased learning – equal for positive and negative outcomes. The *bias* parameter was added to the learning rate for both RL and WM modules.

A recent reanalysis of previous datasets using this task suggests that behavior is better explained by a model combining WM with a tendency to repeat actions regardless of reinforcement history (i.e., habitual learning) (Collins, 2025). Thus, participants do not seem to rely on slow, reinforcement-dependent learning (Collins, 2025) as typically assumed in RL models when performing the RLWM task (Collins et al., 2014; Collins & Frank, 2012; Master et al., 2020). Although we did not preregister it since it was published after our preregistration, we also fitted this alternative model (see Supplementary Materials).

#### GNG Task

The model assigns, for each stimulus (robot), the probability of each action *a_t_* (“go” versus “no-go”) on a trial *t* based on an action weight *W_t_(a, s)* that is passed through a SoftMax function (Sutton & Barto, 2018):

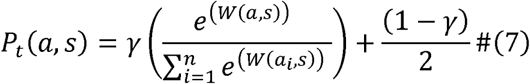

where γ is a lapse rate parameter constrained between 0 and 1.

The initial Q-values for both the go and no-go action are set to 0.5. The action weight consists of the following components:

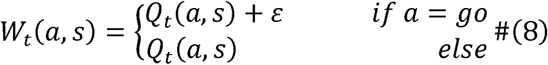

where Q_t_(a,s) is updated as follows:

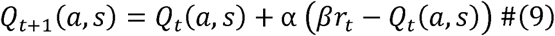

The free parameter α is the learning rate constrained between 0 and 1, and ε ≥ 0 is a free parameter that promotes a tendency to perform a “go” response and represents the motivational or Pavlovian bias. Reinforcements enter the equation through *r_t_* ∈ {0,1}, and β ≥ 0 is a free parameter that determines the effective size of reinforcements.

The β parameter scales the received reward, effectively modulating the perceived impact of outcomes. As such, β controlled reward sensitivity during learning, which is theoretically equivalent to the inverse temperature parameter of the SoftMax function.

#### Effort Task

All three models assume that participants integrate the reward magnitude and the associated physical effort of each option to compute a net subjective value.

The main difference between the models is how the action values are calculated.

### Unconstrained linear model

In the preregistered *unconstrained linear model* (Norbury et al., 2024), the subjective Q-value of the “effort” option (*Q_1_*) was modelled as a reward magnitude scaled by the reward sensitivity parameter minus the effort cost scaled by the effort discounting parameter. The “rest” option (*Q_2_*) was modelled as a reward sensitivity multiplied by 1 which is representing the small, fixed reward:

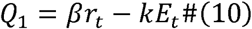

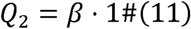

where *r_t_*is the reward magnitude for the effort option displayed to the participant on trial *t*, and *E_t_*is the corresponding effort level displayed to the participants on trial *t*. The free parameter *k* is the physical effort-discounting parameter that captured the degree to which the effort devalues reward where higher *k* indicated greater sensitivity to effort costs. In this preregistered version of the model, *k* is not constrained. The reward sensitivity parameter β controls the impact of reward magnitude and is not constrained either. The higher the β parameter, the higher the impact of reward magnitude on choice probability.

### Constrained linear model

In the second model, *the constrained linear model*, the linear discounting of the effort cost is parametrized as in the preregistered model but the reward sensitivity parameter (*β*) scaled both the reward and the effort term for the “effort” option (*Q_1_*):

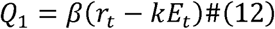

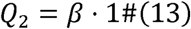

In this model, *β* parameter was constrained such that *β* ≥ 0. The discounting parameter *k* was not constrained as in the previous model.

### Constrained quadratic model

In the third model, *the constrained quadratic model*, effort discounting is parabolic such that the effort cost for a given trial is squared (Lockwood et al., 2022).

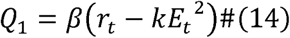

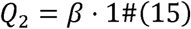

In this model, the *β* parameter is constrained such that *β* ≥ 0 and the *k* parameter is constrained to be between 0 and 1.

For all models, choices are modelled probabilistically using a SoftMax-based choice rule, with an added lapse parameter to account for random responding:

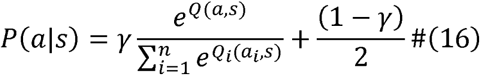

where the free lapse parameter γ serves as an undirected noise parameter that captures occasional random responding or attention lapses during the choice between the two possible options (rest and effort) and is constrained between 0 and 1 in all models.

The winning model is the constrained quadratic model, see Supplemental materials for model comparisons and model recovery results.

#### Parameter and model recovery

We conducted parameter recovery for all winning models finding satisfactory results. The parameters used for the factor analysis were well recovered with r ranging between .68 and .96 (for details see Figure S4, Figure S5 and Figure S7 in Supplementary Materials). Model recovery was carried out for alternative models of the Effort task and RLWM task finding that the predicted behavior of the models could be distinguished. For detailed description of parameter and model recovery, see Supplementary Materials.

#### Choice of the parameters of interest

While the selected computational models would enable us to include more parameters in our analysis, we have pre-selected five parameters of interest. The choice was preregistered and based on the theoretical meaning of the parameters, the results of the pilot study, and the parameter recovery analysis performed after the pilot study. Specifically, we selected the learning rate of the RLWM task, reward sensitivity and motivational bias of the GNG task, foraging exit threshold and effort discounting. We expected that the learning rate and reward sensitivity parameters would underly a RL factor akin to decision acuity (Moutoussis et al., 2021), while exit threshold and effort discounting parameters would underly a separate factor related to vigor. We also hypothesized that the motivational bias would be related to both, because this parameter reflects the tendency to invigorate action in the phase of expected reward at the expense of learning (Guitart-Masip et al., 2014; Oudiette et al., 2019).

### Model-free analysis of performance

To calculate performance levels in each task we used logistic regression and t-tests where appropriate.

### Factor analysis

The exploratory factor analysis (EFA) was estimated by functions *factanal()* and *fa()*. First, we ran the Principal Component Analysis (PCA) on the data to test the hypothesis that 2 factors are meaningful to extract. The number of extracted factors was supported by the two components that had eigenvalues greater than 1 (Kaiser, 1960).

Next, we ran the EFA with principal axis factoring (PA) and varimax rotation. Factor scores were computed using the regression method. The preregistered method to estimate EFA was Maximum Likelihood (ML) method (see Table S6). However, since our parameters were not normally distributed, it is recommended to instead use PA, as it makes no distributional assumptions (Costello & Osborne, 2005; Fabrigar et al., 1999; Watkins, 2018). We therefore report the results of the PA in the main text and the ML for completion in the Supplementary Materials. In fact, simulation studies have shown that PA outperforms ML in datasets with small sample sizes, non-normal distributions, or when the factors are relatively weak (Watkins, 2018).

### Correlation analyses

To investigate associations between the computational parameters of interest and the questionnaire scores, we used Spearman’s rank correlation coefficient (ρ) and applied the correction for multiple comparisons using the Holm method (Holm, 1979).

### Reliability

To estimate test-retest reliability, we calculated the intraclass correlations (ICC, two-way, single measures) between the participants factor scores in Session 1 and Session 2. The obtained ICC indices are interpreted based on the guidelines in the literature (Cicchetti, 1994; Koo & Li, 2016).

### Latent profile analysis (LPA)

To test whether there are distinct underlying profiles in our data based on the multivariate set of parameters used for the FA, we performed an exploratory latent profile analysis (LPA) using the *tidySEM* package version 0.2.11 (Van Lissa, 2019; Van Lissa et al., 2024, 2026). In LPA, the existence of distinct profiles in the data is modelled as a mixture of Gaussians from multiple subpopulations to which participants are probabilistically assigned (Van Lissa et al., 2024). To normalize across different variables, the dataset was standardized by converting the parameter values to z-scores. Using the *mx_model*() function, we estimated models with one to six latent profiles. To test alternative model specifications, variances were set to be either equal or freely estimated across profiles, while covariances were specified as zero, equal, or freely estimated. To ensure that each model contains profiles that can be meaningfully interpreted, all models with profiles including less than 10% of the sample were excluded from the analysis. The best model was selected based on the lowest BIC.

To characterize internal validity and identify how specific parameters contributed to defining the distinct characteristics of each profile, we assessed the differences in the weight patterns of each parameter across profiles. Specifically, we assessed how parameters within profiles differ significantly from the sample means using the built-in function of the tidySEM package which uses Wald z-test. Despite having non-normal distribution for some parameters, the authors of the package claim that they found no evidence in the literature that a more robust measure should be used (Van Lissa et al., 2024). We also compared the parameter values across profiles and considered that profiles differed in a given parameter if the 95% confidence interval did not overlap.

Finally, we explored the relationship between the profiles and the external variables (AMI, ASRM, and CES-D questionnaires). To do so, we used the Bolck–Croon– Hagenaars (*BCH*) method (Bolck et al., 2004), introducing one variable at a time to the model. This method employs a three-step estimation process in which, first, the latent profiles and their parameters are estimated; next, the weights representing the likelihood for an individual to be assigned to a given profile are calculated; and finally, based on these calculations, the mean and confidence interval for the external variable are estimated for each profile (Bolck et al., 2004). The significance testing of the differences among profiles in external variables was done by a likelihood ratio test (LRT).

## Results

The results section contains model-free analysis of behavioral data and computational modelling in accordance with the preregistered hypotheses. We also report a non-preregistered exploratory LPA. The EFA results of a pilot study using a longer version of the battery that were the basis of preregistration are reported in the Supplementary Materials.

### Model-free analysis

To demonstrate that participants choices in the Effort Task are influenced both by effort level and the reward magnitude, we ran a logistic regression showing that the odds were 37% lower for choosing the effort option for each unit increase in effort level (OR = 0.63, 95% CI [0.58, 0.68], *p* < .001). Moreover, the odds were 16% higher for choosing the effortful task for each unit increase in reward magnitude (OR = 1.16, 95% CI [1.11, 1.22], *p* < .001). These effects of reward and effort on choice suggest that increasing effort was discouraging participants from choosing it, while increasing reward magnitude was motivating them. The interaction of effort level and reward magnitude was not significant (OR = 0.99, 95% CI [0.98, 1.00], *p* = .181) suggesting that the effects of effort and reward on choice were independent (see Figure 2A).

To show whether the two conditions in the foraging task were sufficiently sensitive, we compared the exit thresholds for both conditions. There was a significant difference of travel times (*t*(161) = 6.93, *d* = .36, *p* < .001) between the conditions, suggesting that participants adapted their average stay time to adjust for the different travel times (see Figure 2B).

To quantify how participants performed in the RLWM task, we analyzed how learning was affected by each additional stimulus exposure, and whether the speed of learning was different for each set size. For each unit increase in set size, the odds of a correct response decreased by 6.9% (OR = 0.93, 95% CI [0.90, 0.96], *p < .001*), reflecting the increasing difficulty with increasing set size. Furthermore, for each unit increase in stimulus exposure, the odds of a correct response increased by 85% (OR = 1.85, 95% CI [1.79, 1.91], *p < .001*), reflecting overall learning, and lastly, the significant interactions between the set size and stimulus exposure (OR = 0.96, 95% CI [0.95, 0.96], *p < .001*), suggesting that participants were slower to learn in larger set sizes (see Figure 2C).

Finally, in the GNG task, we analyzed how participants learned to respond correctly with each stimulus exposure, and whether learning was affected by the Pavlovian bias. As such the odds were significantly higher for responding correctly for “go” action compared to the “no-go” action (OR = 4.22, 95% CI [3.76, 4.75], *p* < .001), reflecting the general tendency to approach rewarding stimuli rather than abstaining from responding to them. Each additional stimulus exposure increased the odds of a correct response by 14% (OR = 1.14, 95% CI [1.12, 1.15], *p* < .001), reflecting overall learning. Lastly, there was a positive interaction effect of action and stimulus exposure (OR = 0.92, 95% CI [0.90, 0.93], *p* < .001), suggesting that the differences in performance between each stimulus exposure were smaller for the “go” condition. This is because “go” stimuli were generally easier to learn, and therefore the learning curve was not as steep (see Figure 2D).

**Figure 2.**
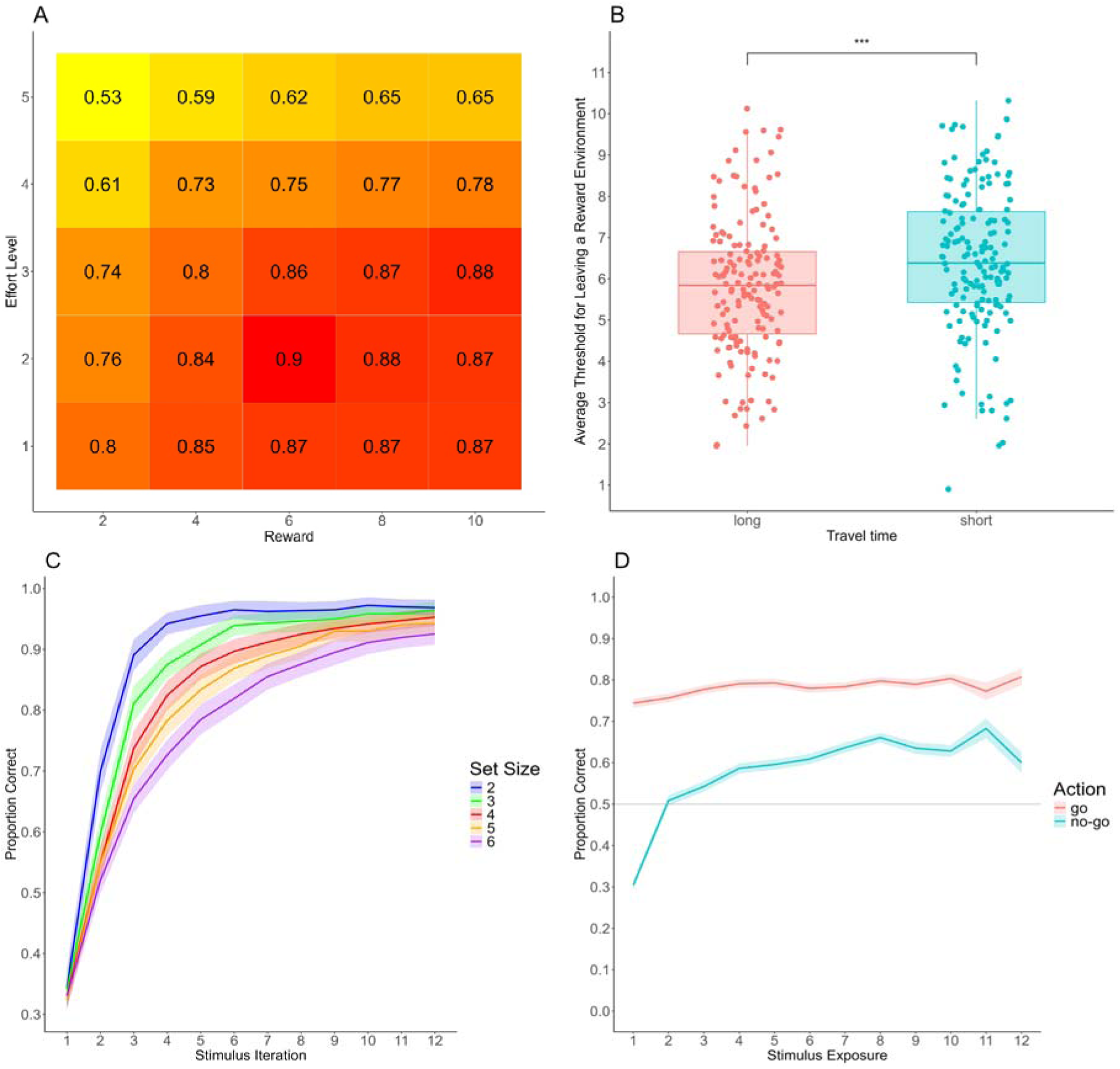
Model-free results of the 4 tasks. *Note.* Results of the model-free analyses from each of the task in the battery. (A) Effort Task: the proportion of accepted effort offers relative all offers averaged across the trials and participants, (B) Foraging task: exit threshold quantifies *reward level at which* participants leave the planet to a new one averaged across the trials for long and short travel times. Each dot represents an individual participant, (C) RLWM Task: Learning curves for each set size averaged across participants representing the proportion of the correct response collapsed across the number of same-stimulus iteration (presentation). Each colour depicts a different set size (memory load), (D) GNG Task: Learning curves averaged across participants representing the proportion of correct responses collapsed across same-stimulus exposure for go and no-go condition.

### Factor analysis: 2-factor solution was supported by the PCA

To evaluate whether RL and Vigor are two underlying constructs that can be quantified using our battery, we conducted an EFA with the 5 parameters of interest: α (learning rate) from the RLWM task, β (reward sensitivity) and ε (motivational bias) from the GNG, Foraging Exit Threshold, and Effort κ (physical effort discounting). Boxplot distributions of the parameters used for the factor analysis are shown in Figure 3 (see Table S4 in Supplementary Materials for the descriptive statistics of all parameters).

**Figure 3.**
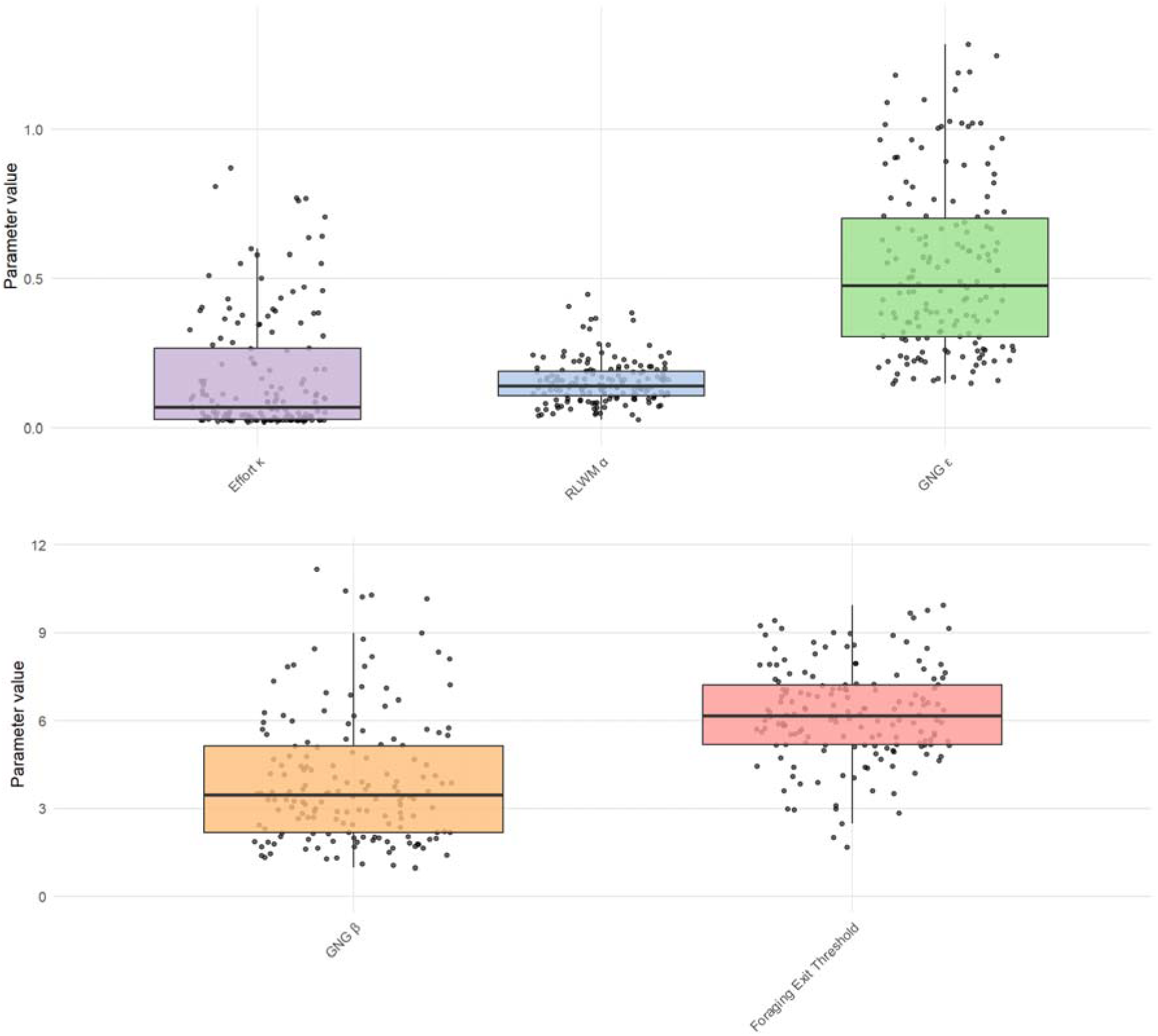
Distribution of the parameters in Session 1.

The PCA revealed that the first two components have eigenvalues > 1, which supported the preregistered hypothesized 2-factor solution (Kaiser, 1960). The EFA, with the 5 parameters of interest, showed the expected pattern of loadings (see Figure 4 or Table S5 in the Supplemental materials for full details).

The EFA showed that RLWM α and GNG β loaded positively on the first factor, explaining 15% of the variance, whereas GNG ε loaded negatively on that factor. We interpreted this factor as reflecting RL, since a high learning rate (α) and reward sensitivity (β) combined with low motivational bias (ε) supported strong performance in RL tasks. We chose GNG β instead of GNG α for the current analysis because the former had a better recoverability profile in our pilot analysis and loaded more strongly on the RL (see the results of our pilot study in the Supplementary Materials). Although we did not preregister the use of the newest model for the RLWM task (Collins, 2025), we tested replacing the learning rate of the RLWM task with WM reliance (WMH ρ) together with the tendency to repeat previous actions (WMH α) from this new model. We show that these two parameters supporting learning in the RLWM task, load together with the reward sensitivity (β) of the GNG task (see Table S11 in Supplementary Materials).

In contrast, GNG ε and Foraging exit threshold loaded positively on a second factor, explaining 5% of the variance, while Effort κ loaded negatively on that factor. We termed this factor vigor, as the factor likely reflects opportunity cost, motivation, and the propensity to pursue rewards. We hypothesized that the respective loadings would be larger than .30. This was supported by the EFA results except that the loading of GNG ε on factor 2 was only .20, albeit the loading was in the hypothesized direction. For a pairwise correlation matrix between the parameters, see Figure S2 in the Supplementary Materials. The factor structure was robust against the removal of univariate and multivariate outliers (see details in the Supplementary Materials).

**Figure 4.**
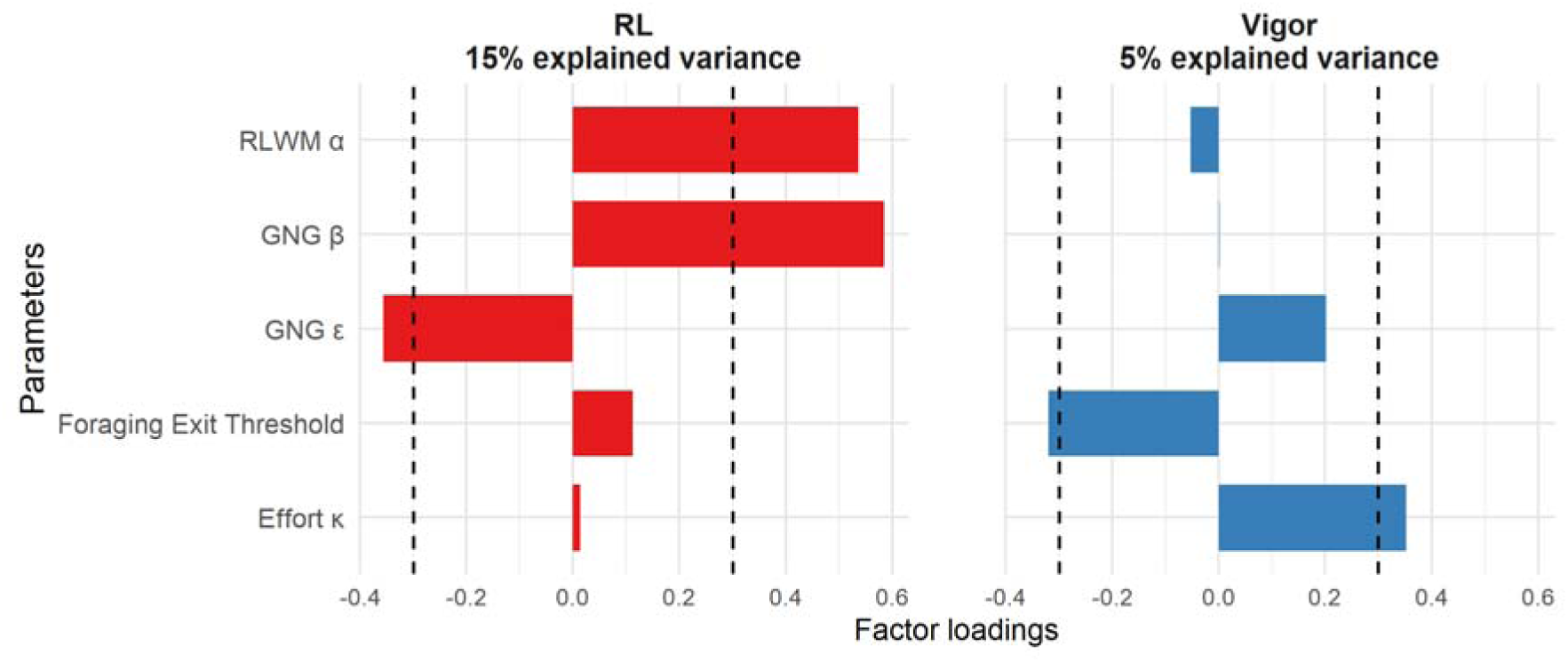
Factor Loadings for Session 1. *Note*. Factor loadings for the two-factor solution including the parameters of interest. Factor 1 was termed RL, while factor 2 was termed vigor. Dashed vertical lines represent the expected hypothesized loadings of -.3 and .3 for the relevant parameters.

In the independent EFA on Session 2, the factor structure from Session 1 was not reproduced for factor 2 which could be no longer interpreted as vigor because *Effort* κ had a positive loading (.18) along with a positive loading of the foraging threshold (.40). Moreover, in factor 1, the negative loading of GNG ε was only -.15 (see Table S7 in Supplementary Materials).

One possible reason for the lack of replication of the factor structure in Session 2 is a smaller sample size compared to Session 1. To explore this possibility, we re-ran the EFA for Session 1 only with participants who had completed Session 2. The factor structure was not preserved, and factors could not be interpreted as RL and Vigor, regardless of extraction method (see Table S12 in Supplementary Materials). This suggests that the sample size in Session 2 may be too small to reliably assess the factor structure.

Assuming the same factor structure across sessions, we observed moderately strong reliability for both factors (factor 1: ICC(2,1) = .65, 95% CI [.48, .77]; factor 2: ICC(2,1) = .67, 95% CI [.51, .78]). In this analysis, the factor scores in Session 2 were estimated using the model parameters estimated in Session 2 and the factor loadings from Session 1 using Thurstone regression approach (Grice, 2001). The obtained ICC indices are considered moderate or good. However, both indices were below the hypothesized threshold of .75. The reliability of individual parameters used in the EFA ranged from .20 to .76, which indicates varying reliability from poor to good (see Reliability Table S13 in Supplementary Materials.

### Predictive validity of RL and Vigor factors

To assess whether the factor scores are associated with symptomatology, we correlated the factor scores with the questionnaire scores. As preregistered based on our pilot study, we found that mania symptoms were negatively associated with factor 1 (ρ = -.30, *p*_holm_ = .002) and positively associated with factor 2 (ρ = .23, *p*_holm_ = .036). Apathy in the behavioral activation (BA) subscale was not significantly associated with factor 1 (ρ = .19, *p*_holm_ = .116), nor with factor 2 (ρ = -.09, *p*_holm_ = 1.000). Depressive symptoms were not associated with either factor 1 (ρ = .00, *p*_holm_ = 1.000) or factor 2 (ρ = .00, *p*_holm_ = 1.000). Next, we ran a series of regression analyses to account for age, sex, and both RL and vigor in the same model. We found that only RL predicted the mania symptoms, *t*(156) = -4.17, β = -1.90, *p* < .001 and not vigor, *t*(156) = 1.63, β = 1.13, *p* = .105. To further test the claim that the RL factor reflects general RL ability, we correlated the factor scores with the obtained performance bonus from the RLWM (ρ = .59, *p*_holm_ < .001), GNG (ρ = .70, *p*_holm_ < .001), and Foraging tasks (ρ = .28, *p*_holm_ = .003). These associations showed that the RL factor reflects the general propensity to make advantageous decisions beyond performance on the RL tasks. The lack of correlation with total rewards in the effort task is expected as the effort task measures subjective preferences and has not optimal policy. On the other hand, vigor was only associated with the obtained performance bonus in the Effort task (ρ = .42, *p*_holm_ < .001), which is not surprising because exerting more effort led to obtaining a higher performance bonus only in that task.

### Exploratory Latent Profile Analysis identified three distinct profiles of participants

We tested alternative models of 1 to 6 profiles with each model having either equal or free variance and zero, equal and free covariance. Given our five parameters of interest, the model with the lowest BIC in the LPA analysis was the model with 3 profiles with free variance and zero covariance (FreeVar3). The FreeVar3 model showed a high entropy (.78), suggesting a good separability of profiles. Further, a high minimum probability of assignment to a profile (.85) points to perfect classification reliability. Finally, the smallest profile accounted for 30.90% of the total sample as such that even the smallest profile was not constituted by only a few participants (see Table S2 for all analyzed models).

The p-values reported below describe which parameter had a significantly different value compared to the sample mean in each profile. The results are summarized in Figure 5. For detailed results, see Table S3 in Supplementary Materials. Profile 1 was composed of 61.24 individuals (38% of the total sample) and was the largest of the profiles. Compared to the sample mean, this profile was defined by lower RLWM α (M = -0.24, SD = 0.12, *p* = .040), lower GNG β (M = -0.42, SD = 0.09, *p* < .001), and lower Effort κ (M = -0.70, SE = 0.01, *p* < .001). Profile 1 showed a non-significant tendency for higher-than-average GNG ε (M = 0.28, SE = 0.15, *p* = .060). Finally, Foraging Exit Threshold did not significantly differ from the mean of the whole sample (M = 0.17, SE = 0.16, *p* = .290). However, Profile 1 showed the highest average Exit Threshold among all profiles, which together with low Effort κ and high GNG ε, suggests that this profile is characterized by high vigor. The conjunction of low RLWM α and GNG β together with high GNG ε also suggests that this profile encompasses participants with low RL ability. As such, we named Profile 1 the low RL / high vigor profile.

Profile 2 was the smallest profile with 48.67 individuals (30% of the total sample). Compared to the sample mean, this profile was defined by lower RLWM α (M = -0.29, SD = 0.11, *p* = .010), lower GNG β (M = -0.45, SD = 0.97, *p* < .001), and higher Effort κ (M = 0.80, SE = 0.20, *p* < .001). Finally, GNG ε (M = 0.14, SE = 0.17, *p* = .420) and Foraging Exit Threshold (M = -0.19, SE = 0.14, *p* = .170) did not significantly differ from the mean of the whole sample. The conjunction of low RLWM α and GNG β together suggests that this profile encompasses participants with low RL ability. Although Foraging Exit Threshold did not significantly differ from the average of the whole sample, Profile 2 showed the lowest average Foraging Exit Threshold among all profiles, which together with high Effort κ, suggest that this profile is characterized by low vigor. As such, we named Profile 2 the low RL / low vigor profile.

Profile 3 was composed of 52.08 individuals (32% of the total sample). Members of this profile displayed higher RLWM α (M = 0.56, SD = 0.20, *p* = .010), higher GNG β (M = 0.90, SE = 0.21, *p* < .001) and lower GNG ε (M = -0.47, SE = 0.10, *p* < .001) when compared to the sample mean. The conjunction of these three parameters suggests that this profile encompasses participants with high RL ability. On the other hand, Profile 3 showed Foraging Exit Threshold (M = -0.02, SE = 0.12, *p* = .840) and Effort κ (M = 0.08, SE = 0.13, *p* = .560) close to the sample mean, suggesting that this profile encompasses participants showing average vigor. As such, we named Profile 3 the high RL / normal vigor profile.

The finding of high apathy in the profile with the highest RL was unexpected. To better understand it, we carried out an exploratory correlation analysis to assess which specific BA items are associated with the RL factor. We found that the highest correlation of BA and RL is driven by items 10. “I don’t like to laze around”, ρ = .19; *p =* .014; *p_holm_* = .083, and 15. “When I have something I need to do, I do it straightaway, so it is out of the way.”, ρ = .17; *p* = .030; *p_holm_*= .148). These exploratory results suggest that the association between RL and apathy might be driven by items of the apathy scale capturing a tendency to deliberate when making decisions.

**Figure 5.**
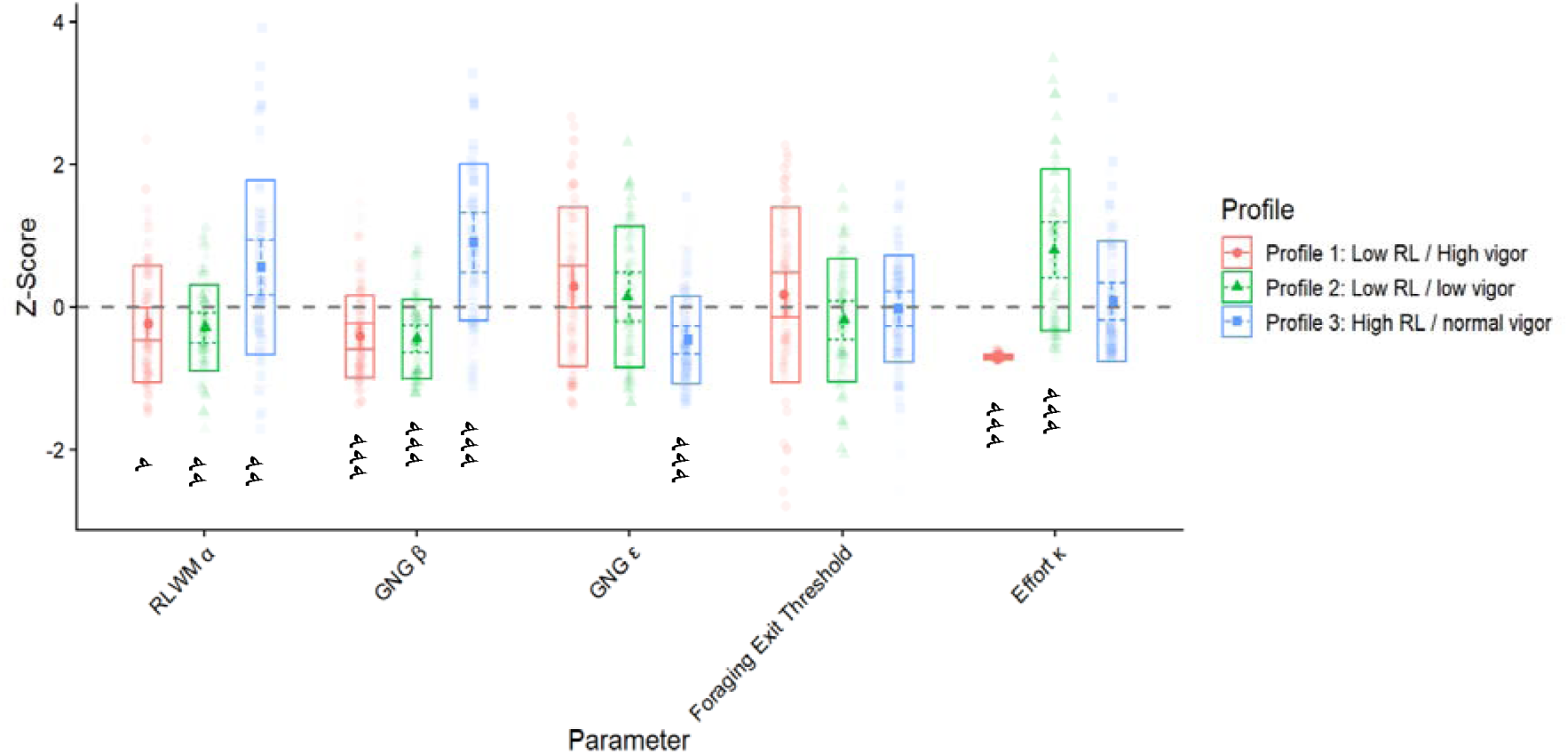
Profile plot of the three profiles from FreeVar 3 profiles model. *Note.* Values on y-axis represent a z-scored parameter value. Each transparent dot represents a participant being assigned to the corresponding profile. The opaque dot represents the parameter mean for the given profile. The whiskers represent a 95% confidence interval. If two intervals within the same parameter do not overlap, then the differences between those two profiles are significantly (p < .050) different from each other. Asterisks indicate the level of statistical significance from the sample mean: *** p < .001; ** p < .01; * p < .05.

Finally, we assessed the relationship between profile membership and symptoms by assessing differences in questionnaire data across profiles using the BCH method. Concerning mania symptoms, significant differences emerged in all three pairwise comparisons (*p <* .019). The low RL / high vigor profile showed higher mania scores (M = 0.43; SE = 0.12), whereas the low RL / low vigor profile exhibited scores close to the sample mean (M = −0.02; SE = 0.14). The high RL / normal vigor profile showed lower levels (M = −0.48; SE = 0.11).

We also found significant differences between the profiles on apathy. The high RL / normal vigor profile differed in apathy from the first (*p* < .001) and the second profile (*p* = .045), respectively. There were no differences between the low RL / high vigor profile and the low RL / low vigor profile (*p* = .059). The low RL / high vigor profile had the lowest levels of BA (M = -0.34, SE = 0.11), while the high RL / normal vigor profile displayed levels close to the sample mean (M = -0.00, SE = 0.14). The high RL / normal vigor exhibited the highest levels of apathy (M = 0.40, SE = 0.14).

No significant pairwise differences emerged between profiles for the depression scores (*p* > .280). All profiles exhibited mean scores close to the overall sample mean: the low RL / high vigor profile (M = 0.05, SE = 0.15), the low RL / low vigor profile (M = −0.11, SE = 0.12) and the high RL / normal vigor profile (M = 0.09, SE = 0.14).

## Discussion

Using our newly developed decision-making battery and factor analysis, we identified the hypothesized factors corresponding to RL and vigor. Factor scores were reliable across the one-to-two-week interval and robust to outlier removal. Moreover, mania symptoms correlated negatively with RL and positively vigor, based on our expectations. However, the association between mania symptoms and vigor was no longer significant after adjusting for RL in post-hoc regression analyses. Furthermore, our results do not suggest relationship between RL or vigor and depression symptoms. Finally, exploratory analyses revealed three distinct profiles in our sample with varying levels of vigor and RL and clear differences in the expression of mania symptoms. Our results suggest the existence of RL and vigor as two aspects underlying decision-making, but while RL has a clear and robust association with mania symptoms, vigor may not predict mania symptoms independently.

Our findings support the notion of RL and vigor as two underlying factors in decision-making. In the tasks selected to capture RL, participants are required to learn stimulus-action-outcome associations (Collins & Frank, 2012; Guitart-Masip et al., 2012; Lawrance et al., 2022; Westbrook et al., 2025; Zorowitz et al., 2025). The GNG β parameter, which reflects the sensitivity to rewards of the instrumental learning system, strongly contributes to the RL factor. In the case of the RLWM task, its model allows us to dissociate slow learning as supported by RL mechanisms from fast, but capacity-limited, learning supported by WM (Collins et al., 2014; Collins & Frank, 2012; Master et al., 2020). In the model, learning ability is captured by the joint effect of the learning rate and WM reliance. However, because WM reliance was not well recovered in our pilots, we only preregistered the inclusion of learning rate in the EFA. Thus, regardless of the exact computational mechanism supporting learning, our RL factor reflects the ability to use outcomes to learn the action contingencies in the RLWM and GNG task. The correlation of the RL factor with performance in the RLWM, GNG and Foraging tasks suggests that the RL factor may represent a general decision-making ability, similar to the decision acuity previously described by Moutoussis et al. (2021). On the other hand, the GNG ε, reflecting the motivational bias, contributes negatively to RL and positively to vigor. This supports the notion that motivational bias impairs learning ability but also contribute to a separate factor that we interpret as vigor.

To quantify vigor in our battery, we chose a physical effort discounting task and a foraging task. In the physical effort discounting task, participants choose between a low reward/low effort offer and a higher reward/higher effort option with varying levels of physical effort and reward magnitude. This manipulation allows a quantification of how much participants discount effort when computing the value of the offers (Bustamante et al., 2024; Chong et al., 2015; Draper et al., 2018; Hewitt et al., 2025; Le Bouc et al., 2016; Le Heron et al., 2018; Lockwood et al., 2017, 2021, 2022; Norbury et al., 2024; Treadway et al., 2009; Westbrook et al., 2020). Our results show that participants’ decisions reflect a tradeoff between reward and number of buttons to be pressed akin to that observed when effort is operationalized as hand gripping (e.g., Lockwood et al., 2017). Our EFA shows that some of the variance in Foraging Exit Threshold covaries with effort discounting and motivational biases. Despite differences between effort and foraging tasks, both tasks impose a cost for obtaining a higher reward and measure the subjective perception of it. Vigor is the policy that optimally balances these costs against expected reward (Niv et al., 2007). Our results suggest that the inverse of effort discounting and patch leaving exit thresholds share a latent factor representing vigor or motivational drive.

Whereas the RL factor explained modest variance and relatively high loadings, the vigor factor had low explained variance and low loadings. These results limit the interpretation of the vigor factor. Even though the existence of the vigor factor is supported by theoretical and experimental work showing that dopamine distinctly modulate RL and vigor (Chong et al., 2015; Dunlop & Nemeroff, 2007; Ianni et al., 2023; Le Heron et al., 2018; Salamone et al., 2007; Westbrook et al., 2020, 2025), the distinction between vigor and RL may be more difficult to capture. Furthermore, we replicated the vigor factor from our pilot study, albeit in the pilot study the vigor factor explained more variance (see the results of our pilot study in the Supplementary Materials). These limitations might be overcome by adding more vigor tasks or adjusting the existing ones. Specifically, participants exhibited a high level of acceptance of the effortful trials in the effort task. Despite having a titration round to find the subjective maximum, a substantial number of participants chose the effort option most of the time. Indeed, 31% of the sample preferred the effort option in more than 90% of the offers. One possible explanation for this high level of acceptance of the effort option is that participants may have found the effort condition more engaging compared to the rest condition. Another potential reason could be a programming error in our code that shortened the training, resulting in participants not experiencing all levels of effort before the actual task started, which was initially planned and tested in our pilot studies.

To provide a meaningful measure of test-retest reliability, we computed factor scores for Session 2 using the loadings in Session 1 (Grice, 2001; Moutoussis et al., 2021). This method is akin to measurement invariance suggested in the literature (Gorsuch, 1988; Meredith, 1993). Factor scores from both sessions exhibited moderately strong test-retest reliability indices. The failure to replicate the factor structure in Session 2 may stem from a smaller sample size with insufficient power in Session 2. Indeed, when we carried out the EFA in Session 1 with only participants who joined both sessions, we did not observe the expected factor structure in Session 1 either. Although we cannot confidently state that the factor structure would be preserved with more participants, these results show that, assuming the same underlying constructs in both sessions, task performance similarly contributes to the factors on both occasions.

To examine predictive validity of the RL and vigor factors, we assessed their relationship with symptoms of mania, apathy, and depression. Mania symptoms correlated negatively with RL and positively with vigor. Previous studies have shown impaired reward learning in a probabilistic reward task among both euthymic and symptomatic patients with bipolar disorder (Pizzagalli et al., 2008). Moreover, higher willingness to exert physical effort in patients with remitted bipolar disorder coincided with their symptom severity (Johnson et al., 2017). These findings align well with our results. However, when including both RL and vigor in a single regression model, only RL predicts mania symptoms, suggesting shared variance between the factors. A similar pattern was revealed by the LPA analysis, showing that the *high RL / normal vigor* profile exhibited lower levels of mania, while the *low RL / high vigor* profile exhibited higher levels of mania. In addition, the *low RL / low vigor* profile exhibited mania levels close to the sample mean despite low levels of vigor. Taken together, our results suggest that low RL may be related to higher levels of mania while vigor does not predict mania symptoms on its own and may not be fully independent from RL.

Contrary to our expectations, we did not find a significant association between vigor and apathy. This is surprising given the association between apathy and higher effort discounting (Bonnelle et al., 2015, 2016; Hartmann et al., 2015; Lockwood et al., 2017) and reduced incentivization by reward (Le Heron et al., 2018). In fact, we observed a weak positive correlation between Effort κ and apathy that did not survive the correction for multiple comparisons (see Figure S14 in Supplementary Materials). The lack of association between apathy and the vigor factor may be due to the small variance attributed to the vigor factor.

Although there was no direct significant association between the RL factor and apathy, the LPA analysis suggested that participants assigned to the *high RL / normal vigor* profile also exhibited higher levels of apathy as compared to participants assigned to a low RL profile regardless of vigor. This suggests that RL ability, and not vigor, as captured by our tasks, may be related to apathy although only in some participants. The higher apathy levels in participants with high levels of RL were not expected and stand in contrast with previous studies showing an association between apathy and reduced learning rates (Yan et al., 2025). The AMI questionnaire used here to assess apathy has commonly been associated with measures of vigor such as foraging (Bustamante et al., 2024; Nair et al., 2023; Trier et al., 2025), effort discounting (Bustamante et al., 2024; Hewitt et al., 2025; Lockwood et al., 2017; Norbury et al., 2024) and impulsivity (Petitet et al., 2022). However, the AMI questionnaire has not been used in previous studies assessing RL abilities. One possible explanation for higher apathy levels in the high RL profile may be that higher levels of perceived apathy in a non-psychiatric sample promote good performance in our tasks. The AMI questionnaire quantifies apathy with items assessing a decrease in goal-oriented behavior but also determination (e.g., “I make decisions firmly and without hesitation). People who perceive themselves deliberating longer or taking more time and effort to decide may score higher on apathy. Indeed, the strong contribution of items assessing urgency or deliberation to the association with the RL factor may provide a possible explanation of the positive relationship between apathy and RL in our data. Future studies will need to determine the generalizability and extent of the association between RL and apathy and anhedonia.

Lastly, our results showed that the depression scores were not associated with either factor, and that the participants exhibited scores close to the sample mean in all profiles of the LPA. Based on the results of our pilot studies, we expected no association between depression and our decision-making factors. Nevertheless, past research has shown that depression was negatively associated with vigor (Ang et al., 2023; Bustamante et al., 2024; Kuhn et al., 2025; Treadway et al., 2009, 2012) and sensitivity to rewards (Costello et al., 2025; Huys et al., 2013; Rupprechter et al., 2018). Our null result provides a challenge for future research, raising the possibility that the effect may be too small to be captured in a general non-clinical sample. This is also in line with the past research in which the decision-acuity factor showed no relationship with a mood-specific factor (Moutoussis et al., 2021).

Some limitations need to be noted. Although online studies allow fast recruitment of a diverse sample, they are less controlled than lab experiments as testing environment and devices differ between participants. Furthermore, online testing imposes constraints on how physical effort can be operationalized. We chose to change the hand gripping effort manipulation to a button pressing manipulation as used in previous studies (Bustamante et al., 2024; Hewitt et al., 2025; Norbury et al., 2024). Although we included gamification elements to increase engagement, we cannot provide evidence that such elements resulted in increased engagement or performance. Instead, we may have inadvertently introduced confounding variables or distracted the participants in unpredictable ways. Finally, despite having sufficient power to detect correlation between the factors and questionnaire scores, our sample size was insufficient to carry out confirmatory factor analysis. Future studies need to elucidate causal relationships of RL and vigor and their relation to dopaminergic processing.

Our study extended the previous research work by constructing a decision-making battery involving both RL and vigor tasks. Using computational modelling and factor analysis, we found preliminary evidence for separate RL and vigor components of decision-making, with RL reflecting a general decision-making ability and vigor representing motivational dimension. RL was negatively related to mania symptoms while vigor was related positively to mania symptoms, although vigor needs further validation. We showed that our online design permits relatively easy and quick administration of the battery and paves the way for the characterization of RL and vigor in patient populations in longitudinal settings.

## Supporting information

Attention checks

Computational modelling

LPA

Outliers

Questionnaires

Regression analyses

Sample

Tasks

## Acknowledgements

This work was supported by two grants awarded by the Swedish Research Council (vetenskapsrådet) to M.G.M. (VR 2021-02046; VR 2024-03094). Parts of the computational analyses were run on the national cluster Tetralith/NSC, enabled by resources provided by the National Academic Infrastructure for Supercomputing in Sweden (NAISS), partially funded by the Swedish Research Council through grant agreement no. 2022-06725.

## Data and code availability

All behavioral data have been deposited at Zenodo and are publicly available at https://doi.org/10.5281/zenodo.23387721. The code for the analyses presented in this paper is openly accessible in a GitHub repository: https://github.com/stepanwenke/Decision-making-and-vigor/.

## Notes

https://doi.org/10.5281/zenodo.23387721

https://github.com/stepanwenke/Decision-making-and-vigor/

### Competing Interest Statement

The authors have declared no competing interest.

### Summary of Updates

- The references to figures and tables in Supplementary Materials were broken. I unlinked the cross references and re-uploaded again the main manuscript file - There was a file called and referenced as Figure S19 but this was a typo and should have been Figure S15 (as in this revision)

## References

Addicott, M. A., Pearson, J. M., Sweitzer, M. M., Barack, D. L., & Platt, M. L. (2017). A Primer on Foraging and the Explore/Exploit Trade-Off for Psychiatry Research. Neuropsychopharmacology, 42(10), 1931–1939. 10.1038/npp.2017.108

Albert, D. A., & Smilek, D. (2023). Comparing attentional disengagement between Prolific and MTurk samples. Scientific Reports, 13(1), 1–13. 10.1038/s41598-023-46048-5

Alexander, L. F., Oliver, A., Burdine, L. K., Tang, Y., & Dunlop, B. W. (2017). Reported maladaptive decision-making in unipolar and bipolar depression and its change with treatment. Psychiatry Research, 257(June), 386–392. 10.1016/j.psychres.2017.08.004

Altman, E. G., Hedeker, D., Peterson, J. L., & Davis, J. M. (1997). The altman self-rating Mania scale. Biological Psychiatry, 42(10), 948–955. 10.1016/S0006-3223(96)00548-3

Ang, Y. S., Lockwood, P., Apps, M. A. J., Muhammed, K., & Husain, M. (2017). Distinct subtypes of apathy revealed by the apathy motivation index. PLoS ONE, 12(1), 1–15. 10.1371/journal.pone.0169938

Ang, Y.-S., Gelda, S. E., & Pizzagalli, D. A. (2023). Cognitive effort-based decision-making in major depressive disorder. Psychological Medicine, 53(9), 4228– 4235. 10.1017/S0033291722000964

Bates, D., Maechler, M., Bolker, B. M., & Walker, S. (2015). Fitting linear mixed-effects models using lme4. Journal of Statistical Software 67: 1—48.

Beierholm, U., Guitart-Masip, M., Economides, M., Chowdhury, R., Düzel, E., Dolan, R., & Dayan, P. (2013). Dopamine modulates reward-related vigor. Neuropsychopharmacology, 38(8), 1495–1503. 10.1038/npp.2013.48

Berke, J. D. (2018). What does dopamine mean? 10.1038/s41593-018-0152-y

Bolck, A., Croon, M., & Hagenaars, J. (2004). Estimating Latent Structure Models with Categorical Variables: One-Step Versus Three-Step Estimators. Political Analysis, 12(1), 3–27. 10.1093/PAN/MPH001

Bonnelle, V., Manohar, S., Behrens, T., & Husain, M. (2016). Individual Differences in Premotor Brain Systems Underlie Behavioral Apathy. Cerebral Cortex, 26(2), 807–819. 10.1093/cercor/bhv247

Bonnelle, V., Veromann, K. R., Burnett Heyes, S., Lo Sterzo, E., Manohar, S., & Husain, M. (2015). Characterization of reward and effort mechanisms in apathy. Journal of Physiology-Paris, 109(1–3), 16–26. 10.1016/J.JPHYSPARIS.2014.04.002

Brown, V. M., Chen, J., Gillan, C. M., & Price, R. B. (2020). Improving the Reliability of Computational Analyses: Model-Based Planning and Its Relationship With Compulsivity. Biological Psychiatry: Cognitive Neuroscience and Neuroimaging, 5(6), 601–609. 10.1016/J.BPSC.2019.12.019

Buelow, M. T., Kowalsky, J. M., & Okdie, B. M. (2024). Test–Retest Reliability of Common Behavioral Decision-Making Tasks: A Multi-Sample, Repeated Measures Study. Archives of Clinical Neuropsychology, 39(3), 378–382. 10.1093/arclin/acad091

Bustamante, L. A., Barch, D. M., Solis, J., Oshinowo, T., Grahek, I., Konova, A. B., Daw, N. D., & Cohen, J. D. (2024). Major depression symptom severity associations with willingness to exert effort and patch foraging strategy. Psychological Medicine, 54(15), 4396–4407. 10.1017/S0033291724002691

Bustamante, L. A., Oshinowo, T., Lee, J. R., Tong, E., Burton, A. R., Shenhav, A. S., Cohen, J. D., & Daw, N. D. (2023). Effort Foraging Task reveals positive correlation between individual differences in the cost of cognitive and physical effort in humans. Proceedings of the National Academy of Sciences of the United States of America, 120(50), 1–33. 10.1073/pnas.2221510120

Carver, C. S., & White, T. L. (1994). Behavioral Inhibition, Behavioral Activation, and Affective Responses to Impending Reward and Punishment: The BIS/BAS Scales. In Journal of Personality and Social Psychology (Vol. 67, Issue 2, pp. 319–333).

Champely, S. (2020). pwr: Basic Functions for Power Analysis (p. 1.3–0) [Data set]. 10.32614/CRAN.package.pwr

Charnov, E. L. (1976). Optimal foraging, the marginal value theorem. Theoretical Population Biology, 9(2), 129–136. 10.1016/0040-5809(76)90040-X

Chong, T. T.-J., Bonnelle, V., Manohar, S., Veromann, K.-R., Muhammed, K., Tofaris, G. K., Hu, M., & Husain, M. (2015). Dopamine enhances willingness to exert effort for reward in Parkinson’s disease. Cortex; a Journal Devoted to the Study of the Nervous System and Behavior, 69, 40–46. 10.1016/j.cortex.2015.04.003

Cicchetti, D. V. (1994). Guidelines , Criteria , and Rules of Thumb for Evaluating Normed and. Psychological Assessment, 6(4), 284–290.

Cohen, S., Kamarck, T., & Mermelstein, R. (1983). A global measure of perceived stress. Journal of Health and Social Behavior, 24(4), 385–396. 10.2307/2136404

Collins, A. G. E. (2025). A habit and working memory model as an alternative account of human reward-based learning. Nature Human Behaviour. 10.1038/s41562-025-02340-0

Collins, A. G. E., Brown, J. K., Gold, J. M., Waltz, J. A., & Frank, M. J. (2014). Working memory contributions to reinforcement learning impairments in schizophrenia. Journal of Neuroscience, 34(41), 13747–13756. 10.1523/JNEUROSCI.0989-14.2014

Collins, A. G. E., & Frank, M. J. (2012). How much of reinforcement learning is working memory, not reinforcement learning? A behavioral, computational, and neurogenetic analysis. European Journal of Neuroscience, 35(7), 1024–1035. 10.1111/j.1460-9568.2011.07980.x

Collins, A. G. E., & Frank, M. J. (2014). Opponent actor learning (OpAL): Modeling interactive effects of striatal dopamine on reinforcement learning and choice incentive. Psychological Review, 121(3), 337–366. 10.1037/a0037015

Constantino, S. M., Dalrymple, J., Gilbert, R. W., Varanese, S., Di Rocco, A., & Daw, N. D. (2017). A Neural Mechanism for the Opportunity Cost of Time. 10.1101/173443

Constantino, S. M., & Daw, N. D. (2015). Learning the opportunity cost of time in a patch-foraging task. *Cognitive*, Affective and Behavioral Neuroscience, 15(4), 837–853. 10.3758/s13415-015-0350-y

Costello, A. B., & Osborne, J. W. (2005). Best practices in exploratory factor analysis: Four recommendations for getting the most from your analysis. *Practical Assessment*, Research, and Evaluation, 10(1). 10.7275/JYJ1-4868

Costello, H., Yamamori, Y., Kieslich, K., Murphy, M., Bobyreva, K., Schrag, A.-E., Howard, R., & Roiser, J. P. (2025). Impaired reward sensitivity in Parkinson’s depression is unresponsive to dopamine treatment. Brain, 148(6), 2122–2134. 10.1093/brain/awaf098

Cousins, D. A., Butts, K., & Young, A. H. (2009). The role of dopamine in bipolar disorder. Bipolar Disorders, 11(8), 787–806. 10.1111/J.1399-5618.2009.00760.X

de Boer, L., Axelsson, J., Chowdhury, R., Riklund, K., Dolan, R. J., Nyberg, L., Bäckman, L., & Guitart-Masip, M. (2019). Dorsal striatal dopamine D1 receptor availability predicts an instrumental bias in action learning. Proceedings of the National Academy of Sciences of the United States of America, 116(1), 261–270. 10.1073/pnas.1816704116

de Leeuw, J. R., Gilbert, R. A., & Luchterhandt, B. (2023). jsPsych: Enabling an Open-Source Collaborative Ecosystem of Behavioral Experiments. Journal of Open Source Software, 8(85), 5351. 10.21105/joss.05351

Draper, A., Koch, R. M., Van Der Meer, J. W., Apps, M. A., Pickkers, P., Husain, M., & Van Der Schaaf, M. E. (2018). Effort but not Reward Sensitivity is Altered by Acute Sickness Induced by Experimental Endotoxemia in Humans. Neuropsychopharmacology, 43(5), 1107–1118. 10.1038/npp.2017.231

Dunlop, B. W., & Nemeroff, C. B. (2007). The role of dopamine in the pathophysiology of depression. Archives of General Psychiatry, 64(3), 327– 337. 10.1001/archpsyc.64.3.327

Endrass, T., & Ullsperger, M. (2021). Decision-making as transdiagnostic construct for mental health research. Neuron, 109(12), 1912–1914. 10.1016/j.neuron.2021.05.035

Fabrigar, L. R., MacCallum, R. C., Wegener, D. T., & Strahan, E. J. (1999). Evaluating the use of exploratory factor analysis in psychological research. Psychological Methods, 4(3), 272. 10.1037/1082-989X.4.3.272

Gorsuch, R. L. (1988). Exploratory Factor Analysis.

Granwald, T., Dayan, P., Lengyel, M., & Guitart-Masip, M. (2025). A task-invariant prior explains trial-by-trial active avoidance behaviour across gain and loss tasks. Communications Psychology 2025 3:1, 3(1), 82-. 10.1038/s44271-025-00254-1

Grice, J. W. (2001). Computing and evaluating factor scores. Psychological Methods, 6(4), 430–450. 10.1037/1082-989X.6.4.430

Guitart-Masip, M., Beierholm, U. R., Dolan, R., Duzel, E., & Dayan, P. (2011). Vigor in the face of fluctuating rates of reward: An experimental examination. Journal of Cognitive Neuroscience, 23(12), 3933–3938. 10.1162/jocn_a_00090

Guitart-Masip, M., Duzel, E., Dolan, R., & Dayan, P. (2014). Action versus valence in decision making. Trends in Cognitive Sciences, 18(4), 194–202. 10.1016/j.tics.2014.01.003

Guitart-Masip, M., Huys, Q. J. M., Fuentemilla, L., Dayan, P., Duzel, E., & Dolan, R. J. (2012). Go and no-go learning in reward and punishment: Interactions between affect and effect. NeuroImage, 62(1), 154–166. 10.1016/j.neuroimage.2012.04.024

Hamid, A. A., Pettibone, J. R., Mabrouk, O. S., Hetrick, V. L., Schmidt, R., Vander Weele, C. M., Kennedy, R. T., Aragona, B. J., & Berke, J. D. (2016). Mesolimbic dopamine signals the value of work. 19(1). 10.1038/nn.4173

Harhen, N. C., & Bornstein, A. M. (2023). Overharvesting in human patch foraging reflects rational structure learning and adaptive planning. Proceedings of the National Academy of Sciences of the United States of America, 120(13), e2216524120. 10.1073/pnas.2216524120

Hartmann, M. N., Hager, O. M., Reimann, A. V., Chumbley, J. R., Kirschner, M., Seifritz, E., Tobler, P. N., & Kaiser, S. (2015). Apathy But Not Diminished Expression in Schizophrenia Is Associated With Discounting of Monetary Rewards by Physical Effort. Schizophrenia Bulletin, 41(2), 503–512. 10.1093/schbul/sbu102

Hewitt, S. R. C., Norbury, A., Huys, Q. J. M., & Hauser, T. U. (2025). Day-to-day fluctuations in motivation drive effort-based decision-making. Proceedings of the National Academy of Sciences, 122(12), e2417964122. 10.1073/pnas.2417964122

Holm, S. (1979). A Simple Sequentially Rejective Multiple Test Procedure. Scandinavian Journal of Statistics, 6(2), 65–70.

Hubert, M., & Vandervieren, E. (2008). An adjusted boxplot for skewed distributions. Computational Statistics & Data Analysis, 52(12), 5186–5201. 10.1016/J.CSDA.2007.11.008

Huys, Q. J. M., Cools, R., Gölzer, M., Friedel, E., Heinz, A., Dolan, R. J., & Dayan, P. (2011). Disentangling the roles of approach, activation and valence in instrumental and pavlovian responding. PLoS Computational Biology, 7(4). 10.1371/journal.pcbi.1002028

Huys, Q. J. M., Daw, N. D., & Dayan, P. (2015). Depression: A Decision-Theoretic Analysis. Annual Review of Neuroscience, 38, 1–23. 10.1146/annurev-neuro-071714-033928

Huys, Q., Pizzagalli, D. A., Bogdan, R., & Dayan, P. (2013). Mapping anhedonia onto reinforcement learning: A behavioural meta-analysis. Biology of Mood & Anxiety Disorders, 3(1), 12. 10.1186/2045-5380-3-12

Huys, Q., Vogelstein, J., & Dayan, P. (2008). Psychiatry: Insights into depression through normative decision-making models. Advances in Neural Information Processing Systems, 21. https://proceedings.neurips.cc/paper_files/paper/2008/hash/d04d42cdf14579cd294e5079e0745411-Abstract.html

Hyman, S. E. (2007). Can neuroscience be integrated into the DSM-V? Nature Reviews Neuroscience, 8(9), 725–732. 10.1038/nrn2218

Ianni, A. M., Eisenberg, D. P., Boorman, E. D., Constantino, S. M., Hegarty, C. E., Gregory, M. D., Masdeu, J. C., Kohn, P. D., Behrens, T. E., & Berman, K. F. (2023). PET-measured human dopamine synthesis capacity and receptor availability predict trading rewards and time-costs during foraging. Nature Communications, 14(1), 1–13. 10.1038/s41467-023-41897-0

Johnson, S. L., Swerdlow, B. A., Treadway, M., Tharp, J. A., & Carver, C. S. (2017). Willingness to Expend Effort Toward Reward and Extreme Ambitions in Bipolar I Disorder. Clinical Psychological Science, 5(6), 943–951. 10.1177/2167702617718181

Judd, N., Aristodemou, M., Klingberg, T., & Kievit, R. (2024). Interindividual Differences in Cognitive Variability Are Ubiquitous and Distinct From Mean Performance in a Battery of Eleven Tasks. Journal of Cognition, 7(1), 1–16. 10.5334/joc.371

Kaiser, H. F. (1960). The application of electronic computers to factor analysis. Educational and Psychological Measurement, 20(1), 141–151.

Koo, T. K., & Li, M. Y. (2016). A Guideline of Selecting and Reporting Intraclass Correlation Coefficients for Reliability Research. Journal of Chiropractic Medicine, 15(2), 155–163. 10.1016/j.jcm.2016.02.012

Kuhn, M., Palermo, E. H., Pagnier, G., Blank, J. M., Steinberger, D. C., Long, Y., Nowicki, G., Cooper, J. A., Treadway, M. T., Frank, M. J., & Pizzagalli, D. A. (2025). Computational Phenotyping of Effort-Based Decision Making in Unmedicated Adults With Remitted Depression. Biological Psychiatry: Cognitive Neuroscience and Neuroimaging, 10(6), 607–615. 10.1016/j.bpsc.2025.02.006

Lawrance, E. L., Gagne, C. R., O’Reilly, J. X., Bjisterbosch, J., & Bishop, S. J. (2022). The Computational and Neural Substrates of Ambiguity Avoidance in Anxiety. Computational Psychiatry, 6(1), 8–33. 10.5334/cpsy.67

Lawrence, M. A. (2026). ez: Easy Analysis and Visualization of Factorial Experiments. 10.32614/CRAN.package.ez

Le Bouc, R., Rigoux, L., Schmidt, L., Degos, B., Welter, M.-L., Vidailhet, M., Daunizeau, J., & Pessiglione, M. (2016). Computational Dissection of Dopamine Motor and Motivational Functions in Humans. The Journal of Neuroscience, 36(25), 6623–6633. 10.1523/JNEUROSCI.3078-15.2016

Le Heron, C., Kolling, N., Plant, O., Kienast, A., Janska, R., Ang, Y. S., Fallon, S., Husain, M., & Apps, M. A. J. (2020). Dopamine Modulates Dynamic Decision-Making during Foraging. Journal of Neuroscience, 40(27), 5273–5282. 10.1523/JNEUROSCI.2586-19.2020

Le Heron, C., Plant, O., Manohar, S., Ang, Y.-S., Jackson, M., Lennox, G., Hu, M. T., & Husain, M. (2018). Distinct effects of apathy and dopamine on effort-based decision-making in Parkinson’s disease. Brain, 141(5), 1455–1469. 10.1093/brain/awy110

Lee, D. (2013). Decision Making: From Neuroscience to Psychiatry. Neuron, 78(2), 233–248. 10.1016/j.neuron.2013.04.008

Lenow, J. K., Constantino, S. M., Daw, N. D., & Phelps, E. A. (2017). Chronic and acute stress promote overexploitation in serial decision making. Journal of Neuroscience, 37(23), 5681–5689. 10.1523/JNEUROSCI.3618-16.2017

Lenth, R. V., & Piaskowski, J. (2025). emmeans: Estimated Marginal Means, aka Least-Squares Means. 10.32614/CRAN.package.emmeans

Lloyd, A., Viding, E., McKay, R., & Furl, N. (2023). Understanding patch foraging strategies across development. Trends in Cognitive Sciences, 27(11), 1085– 1098. 10.1016/j.tics.2023.07.004

Lockwood, P. L., Abdurahman, A., Gabay, A. S., Drew, D., Tamm, M., Husain, M., & Apps, M. A. J. (2021). Aging Increases Prosocial Motivation for Effort. Psychological Science, 32(5), 668–681. 10.1177/0956797620975781

Lockwood, P. L., Hamonet, M., Zhang, S. H., Ratnavel, A., Salmony, F. U., Husain, M., & Apps, M. A. J. (2017). Prosocial apathy for helping others when effort is required. Nature Human Behaviour, 1(7), 1–23. 10.1038/s41562-017-0131

Lockwood, P. L., Wittmann, M. K., Nili, H., Matsumoto-Ryan, M., Abdurahman, A., Cutler, J., Husain, M., & Apps, M. A. J. (2022). Distinct neural representations for prosocial and self-benefiting effort. Current Biology, 32(19), 4172–4185.e7. 10.1016/j.cub.2022.08.010

Lüdecke, D., Ben-Shachar, M. S., Patil, I., Wiernik, B. M., Bacher, E., Thériault, R., & Makowski, D. (2022). easystats: Framework for Easy Statistical Modeling, Visualization, and Reporting. CRAN. 10.32614/CRAN.package.easystats

Master, S. L., Eckstein, M. K., Gotlieb, N., Dahl, R., Wilbrecht, L., & Collins, A. G. E. (2020). Distentangling the systems contributing to changes in learning during adolescence. Developmental Cognitive Neuroscience, 41. 10.1016/j.dcn.2019.100732

McClure, S. M., Daw, N. D., & Montague, P. R. (2003). A computational substrate for incentive salience. Trends in Neurosciences, 26(8), 423–428. 10.1016/s0166-2236(03)00177-2

Meredith, W. (1993). Measurement Invariance, Factor Analysis and Factorial Invariance. Psychometrika, 58(4), 525–543. 10.1007/BF02294825

Mkrtchian, A., Valton, V., & Roiser, J. P. (2023). Reliability of Decision-Making and Reinforcement Learning Computational Parameters. Computational Psychiatry, 7(1), 30–46. 10.5334/CPSY.86

Mohebi, A., Pettibone, J. R., Hamid, A. A., Wong, J.-M. T., Vinson, L. T., Patriarchi, T., Tian, L., Kennedy, R. T., & Berke, J. D. (2019). Dissociable dopamine dynamics for learning and motivation. Nature, 570(7759), 65–70. 10.1038/s41586-019-1235-y

Montague, P. R., Dayan, P., & Sejnowski, T. J. (1996). A framework for mesencephalic dopamine systems based on predictive Hebbian learning. Journal of Neuroscience, 16(5), 1936–1947. 10.1523/JNEUROSCI.16-05-01936.1996

Montague, P. R., Dolan, R. J., Friston, K. J., & Dayan, P. (2012). Computational psychiatry. Trends in Cognitive Sciences, 16(1), 72–80. 10.1016/j.tics.2011.11.018

Moutoussis, M., Garzón, B., Neufeld, S., Bach, D. R., Rigoli, F., Goodyer, I., Bullmore, E., Fonagy, P., Jones, P., Hauser, T., Romero-Garcia, R., St Clair, M., Vértes, P., Whitaker, K., Inkster, B., Prabhu, G., Ooi, C., Toseeb, U., Widmer, B., … Dolan, R. J. (2021). Decision-making ability, psychopathology, and brain connectivity. Neuron, 109(12), 2025–2040.e7. 10.1016/j.neuron.2021.04.019

Mukherjee, D., Lee, S., Kazinka, R., D. Satterthwaite, T., & Kable, J. W. (2020). Multiple Facets of Value-Based Decision Making in Major Depressive Disorder. Scientific Reports, 10(1), 3415. 10.1038/s41598-020-60230-z

Murphy, F. C., Rubinsztein, J. S., Michael, A., Rogers, R. D., Robbins, T. W., Paykel, E. S., & Sahakian, B. J. (2001). Decision-making cognition in mania and depression. Psychological Medicine, 31(4), 679–693. 10.1017/S0033291701003804

Nair, A., Niyogi, R. K., Shang, F., Tabrizi, S. J., Rees, G., & Rutledge, R. B. (2023). Opportunity cost determines free-operant action initiation latency and predicts apathy. Psychological Medicine, 53(5), 1850–1859. 10.1017/S0033291721003469

Niv, Y. (2009). Reinforcement learning in the brain. *Journal of Mathematical Psychology*, Special Issue: Dynamic Decision Making, 53(3), 139–154. 10.1016/j.jmp.2008.12.005

Niv, Y., Daw, N. D., Joel, D., & Dayan, P. (2007). Tonic dopamine: Opportunity costs and the control of response vigor. Psychopharmacology, 191(3), 507–520. 10.1007/s00213-006-0502-4

Norbury, A., Hauser, T. U., Fleming, S. M., Dolan, R. J., & Huys, Q. J. M. M. (2024). Different components of cognitive-behavioral therapy affect specific cognitive mechanisms. In Sci. Adv (Vol. 10, Issue 13, p. 3222). 10.1126/sciadv.adk3222

Ooms, J. (2014). The jsonlite Package: A Practical and Consistent Mapping Between JSON Data and R Objects.

Oudiette, D., Vinckier, F., Bioud, E., & Pessiglione, M. (2019). A Pavlovian account for paradoxical effects of motivation on controlling response vigour. Scientific Reports, 9(1), 1–13. 10.1038/s41598-019-43936-7

Parker, A. M., & Fischhoff, B. (2005). Decision-making competence: External validation through an individual-differences approach. Journal of Behavioral Decision Making, 18(1), 1–27. 10.1002/bdm.481

Paulus, M. P., Huys, Q. J. M., & Maia, T. V. (2016). A Roadmap for the Development of Applied Computational Psychiatry. *Biological Psychiatry: Cognitive Neuroscience and Neuroimaging*, Model-Based and Machine-Learning Approaches in Applied Computational Psychiatry, 1(5), 386–392. 10.1016/j.bpsc.2016.05.001

Petitet, P., Zhao, S., Drew, D., Manohar, S. G., & Husain, M. (2022). Dissociable behavioural signatures of co-existing impulsivity and apathy in decision-making. Scientific Reports, 12(1), 1–12. 10.1038/s41598-022-25882-z

Pike, A. C., Board, R., Travers, E., Croal, M., Lam, Y. C., Valton, V., Robinson, O. J., & Roiser, J. P. (2026). The recoverability, reliability, and generalizability of reward processing parameters and relation to mental health symptoms. Psychological Medicine, 56, e238. 10.1017/S0033291726105340

Pike, A. C., Tan, K. H. T., Tromblee, H., Wing, M., & Robinson, O. J. (2024). Test-Retest Reliability of Two Computationally-Characterised Affective Bias Tasks. Computational Psychiatry, 8(1). 10.5334/cpsy.92

Pizzagalli, D. A. (2014). Depression, stress, and anhedonia: Toward a synthesis and integrated model. Annual Review of Clinical Psychology, 10, 393–423. 10.1146/annurev-clinpsy-050212-185606

Pizzagalli, D. A., Goetz, E., Ostacher, M., Iosifescu, D. V., & Perlis, R. H. (2008). Euthymic Patients with Bipolar Disorder Show Decreased Reward Learning in a Probabilistic Reward Task. Biological Psychiatry, 64(2), 162–168. 10.1016/j.biopsych.2007.12.001

Pooseh, S., Bernhardt, N., Guevara, A., Huys, Q. J. M., & Smolka, M. N. (2017). Value-based decision-making battery: A Bayesian adaptive approach to assess impulsive and risky behavior. 10.3758/s13428-017-0866-x

R Core Team. (2025). R: A Language and Environment for Statistical Computing. R Foundation for Statistical Computing. https://www.r-project.org/

Radloff, L. S. (1977). The CES-D Scale: A Self-Report Depression Scale for Research in the General Population. Applied Psychological Measurement, 1(3), 385–401. 10.1177/014662167700100306

Ramírez-Martín, A., Ramos-Martín, J., Mayoral-Cleries, F., Moreno-Küstner, B., & Guzman-Parra, J. (2020). Impulsivity, decision-making and risk-taking behaviour in bipolar disorder: A systematic review and meta-analysis. Psychological Medicine, 50(13), 2141–2153. 10.1017/S0033291720003086

Revelle, W. (2015). Package ’psych’—Procedures for Psychological, Psychometric and Personality Research. In R Package (pp. 1–358). Northwestern University.

Robinson, D., Hayes, A., Couch, S., & Hvitfeldt, E. (2026). broom: Convert Statistical Objects into Tidy Tibbles. 10.32614/CRAN.package.broom

Rupprechter, S., Stankevicius, A., Huys, Q. J. M., Steele, J. D., & Seriès, P. (2018). Major Depression Impairs the Use of Reward Values for Decision-Making. Scientific Reports, 8(1), 1–8. 10.1038/s41598-018-31730-w

Salamone, J. D., & Correa, M. (2002). Motivational views of reinforcement: Implications for understanding the behavioral functions of nucleus accumbens dopamine. Behavioural Brain Research, 137(1–2), 3–25. 10.1016/S0166-4328(02)00282-6

Salamone, J. D., & Correa, M. (2012). The Mysterious Motivational Functions of Mesolimbic Dopamine. Neuron, 76(3), 470–485. 10.1016/j.neuron.2012.10.021

Salamone, J. D., Correa, M., Farrar, A., & Mingote, S. M. (2007). Effort-related functions of nucleus accumbens dopamine and associated forebrain circuits. Psychopharmacology, 191(3), 461–482. 10.1007/s00213-006-0668-9

Satoh, T., Nakai, S., Sato, T., & Kimura, M. (2003). Correlated Coding of Motivation and Outcome of Decision by Dopamine Neurons. Journal of Neuroscience, 23(30).

Schaaf, J. V., Weidinger, L., Molleman, L., & van den Bos, W. (2024). Test–retest reliability of reinforcement learning parameters. Behavior Research Methods, 56(5), 4582–4599. 10.3758/s13428-023-02203-4

Scholl, J., & Klein-Flügge, M. (2018). Understanding psychiatric disorder by capturing ecologically relevant features of learning and decision-making. Behavioural Brain Research, 355(March 2017), 56–75. 10.1016/j.bbr.2017.09.050

Schultz, W. (2016). Dopamine reward prediction error coding. Dialogues in Clinical Neuroscience, 18(1), 23–32. 10.31887/dcns.2016.18.1/wschultz

Schultz, W. (2019). Recent advances in understanding the role of phasic dopamine activity. F1000Research, 8, 1–12. 10.12688/f1000research.19793.1

Schultz, W., Dayan, P., & Montague, P. R. (1997). A neural substrate of prediction and reward. Science, 275(5306), 1593–1599. 10.1126/science.275.5306.1593

Snaith, R. P., Hamilton, M., Morley, S., Humayan, A., Hargreaves, D., & Trigwell, P. (1995). A scale for the assessment of hedonic tone. The Snaith-Hamilton Pleasure Scale. British Journal of Psychiatry, 167(JULY), 99–103. 10.1192/bjp.167.1.99

Stephens, D. W., & Krebs, J. R. (1986). Foraging Theory. Princeton University Press.

Sutton, R. S. ., & Barto, A. G. . (2018). Reinforcement Learning: An Introduction: 2nd edition (Vol. 1, Issue 1). The MIT Press.

Swart, J. C., Froböse, M. I., Cook, J. L., Geurts, D. E., Frank, M. J., Cools, R., & den Ouden, H. E. (2017). Catecholaminergic challenge uncovers distinct Pavlovian and instrumental mechanisms of motivated (in)action. eLife, 6, e22169. 10.7554/eLife.22169

Thériault, R. (2023). {rempsyc}: Convenience functions for psychology. Journal of Open Source Software, 8(87), 5466. 10.21105/joss.05466

Treadway, M. T., Bossaller, N. A., Shelton, R. C., & Zald, D. H. (2012). Effort-based decision-making in major depressive disorder: A translational model of motivational anhedonia. Journal of Abnormal Psychology, 121(3), 553–558. 10.1037/a0028813

Treadway, M. T., Buckholtz, J. W., Schwartzman, A. N., Lambert, W. E., & Zald, D. H. (2009). Worth the “EEfRT”? The effort expenditure for rewards task as an objective measure of motivation and anhedonia. PLoS ONE, 4(8), 1–9. 10.1371/journal.pone.0006598

Treadway, M. T., & Zald, D. H. (2011). Reconsidering anhedonia in depression: Lessons from translational neuroscience. Neuroscience and Biobehavioral Reviews, 35(3), 537–555. 10.1016/j.neubiorev.2010.06.006

Treadway, M. T., & Zald, D. H. (2014). Translational Models of Dopaminergic Mechanisms for Motivational Deficits in Anhedonic Patients. In M. S. Ritsner (Ed.), Anhedonia: A Comprehensive Handbook Volume I: Conceptual Issues And Neurobiological Advances (pp. 107–117). Springer Netherlands. 10.1007/978-94-017-8591-4_7

Trier, H. A., O’Reilly, J. X., Spiering, L., Yishan, S. M., Kolling, N., Rushworth, M. F. S., & Scholl, J. (2025). Emotions and individual differences shape human foraging under threat. Nature Mental Health, 3(4), 444–465. 10.1038/s44220-025-00393-8

Van Lissa, C. J. (2019). *tidySEM* [R]. https://github.com/cjvanlissa/tidySEM

Van Lissa, C. J., Garnier-Villarreal, M., & Anadria, D. (2024). Recommended Practices in Latent Class Analysis Using the Open-Source R-Package tidySEM. Structural Equation Modeling, 31(3), 526–534. 10.1080/10705511.2023.2250920

Van Lissa, C. J. van, Garnier-Villarreal, M., & Gootjes, F. C. (2026). *tidySEM: Tidy Structural Equation Modeling* (Version 0.2.12) [Computer software]. https://cran.r-project.org/web/packages/tidySEM/index.html

Wang, A. Y., Miura, K., & Uchida, N. (2013). The dorsomedial striatum encodes net expected return, critical for energizing performance vigor. Nature Neuroscience, 16(5), 639–647. 10.1038/nn.3377

Watkins, C. J. C. H., & Dayan, P. (1992). Q-learning. Machine Learning, 8(3), 279–292. 10.1007/BF00992698

Watkins, M. W. (2018). Exploratory Factor Analysis: A Guide to Best Practice. Journal of Black Psychology, 44(3), 219–246. 10.1177/0095798418771807

Westbrook, A., van den Bosch, R., Hofmans, L., Papadopetraki, D., Määttä, J. I., Collins, A. G. E., Frank, M. J., & Cools, R. (2025). Striatal dopamine can enhance both fast working memory, and slow reinforcement learning, while reducing implicit effort cost sensitivity. Nature Communications, 16(1). 10.1038/s41467-025-61099-0

Westbrook, A., Van Den Bosch, R., Määttä, J. I., Hofmans, L., Papadopetraki, D., Cools, R., & Frank, M. J. (2020). Dopamine promotes cognitive effort by biasing the benefits versus costs of cognitive work. Science, 367(6484), 1362– 1366. 10.1126/science.aaz5891

Wickham, H. (2016). ggplot2: Elegant Graphics for Data Analysis. Springer-Verlag New York. https://ggplot2.tidyverse.org

Wickham, H., Averick, M., Bryan, J., Chang, W., McGowan, L. D., François, R., Grolemund, G., Hayes, A., Henry, L., Hester, J., Kuhn, M., Pedersen, T. L., Miller, E., Bache, S. M., Müller, K., Ooms, J., Robinson, D., Seidel, D. P., Spinu, V., … Yutani, H. (2019). Welcome to the {tidyverse}. Journal of Open Source Software, 4(43), 1686. 10.21105/joss.01686

Wickham, H., François, R., Henry, L., Müller, K., & Vaughan, D. (2023). dplyr: A Grammar of Data Manipulation. 10.32614/CRAN.package.dplyr

Wickham, H., Vaughan, D., & Girlich, M. (2025). tidyr: Tidy Messy Data. 10.32614/CRAN.package.tidyr

Yan, X., Ebitz, R. B., Grissom, N., Darrow, D. P., & Herman, A. B. (2025). Distinct Computational Mechanisms of Uncertainty Processing Explain Opposing Exploratory Behaviors in Anxiety and Apathy. Biological Psychiatry: Cognitive Neuroscience and Neuroimaging, 10(9), 954–963. 10.1016/j.bpsc.2025.01.005

Zorowitz, S., Karni, G., Paredes, N., Daw, N., & Niv, Y. (2025). Improving the Reliability of the Pavlovian Go/No-Go Task for Computational Psychiatry Research. Computational Psychiatry, 9(1), 231–252. 10.5334/cpsy.127

