## Supplementary material for "Assessing reinforcement learning and vigor with an online decision-making battery": Attention checks

Within tasks, participants were prompted to press a specific key within 10 seconds at different times throughout the tasks. In each questionnaire, there was an extra item asking to select a specific answer (e.g., Choose “Definitely agree” to prove that you paid attention here”). In total, there were 21 attention checks embedded in the tasks and questionnaires.

1 participant failed two attention checks within the GNG task and was therefore not invited to the second session (shown in Figure 1); no other participant failed 2 or more attention checks within one task or within questionnaires for any of the sessions. Further, in Session 1, there was one participant who failed one attention check in the Effort and Foraging tasks and six more participants who failed one attention check each. In Session 2, two participants failed one attention check each.
