## Supplementary material for "Assessing reinforcement learning and vigor with an online decision-making battery": Computational modelling

### Parameter estimation

Parameter estimation followed the approach used in similar previous studies (Guitart-Masip et al., 2012; Huys et al., 2011). Parameters were estimated using an iterative Bayesian hierarchical approach using an expectation-maximization (EM) method initialized with Gaussian priors (mean = 0, variance = 1) on all free parameters. Specifically, the parameter estimation was performed using the *fminsearch* function, and in each iteration, the group-level mean and variance were used as a prior for the next iteration. This iterative process continued until it reached the convergence criterion, defined as the change between the current and last iteration’s group parameters and log-likelihoods being smaller than .01. We used this iterative approach to prevent individual parameters from taking extreme values. Furthermore, to mitigate the risk of converging to local minima, we performed each fitting 10 times with different random initializations. When converged, a sum of the negative penalized log-likelihoods (PL) for each initialization was checked to choose the best fit.

### Parameter recovery

We carried out parameter recovery by assessing the correspondence between the fitted and observed parameters using Pearson correlation (r) and coefficient of determination (R_2_). Specifically, the parameter recovery was done by generating 1000 synthetic datasets by sampling the mean and variance of the estimated parameter values observed in our data and then fitting the model that generated the data to these synthetic datasets using the same hierarchical method as during the parameter estimation. We obtained good to excellent recoveries for the parameters of interest used in the current study for factor analysis and LPA: RLWM alpha = .68, GNG β = .81, GNG epsilon = .86 and Effort κ = .96- See the scatter plots below of all parameters below with the main EFA parameters highlighted.

### Model comparison

We employed a computational model used previously for a GNG task (Guitart-Masip et al., 2012). As such, no model comparisons were made for this model.

For the Effort task, in addition to the preregistered model, we compared two alternative models described in the Computational Modelling section to ensure that we use the best model for this newly developed task in our lab (for the details see Supplemental materials).

For the RLWM task, we used an established RLWM model (Master et al., 2020). However, more recently, Collins (2025) demonstrated that behavior in the task may be better captured by a model that only incorporates working memory and habitual learning (WMH). We performed parameter recovery for this model and carried out a model comparison between the WMH and RLWM models. Because the WMH and the RLWM have the same number of parameters, we compared the two models directly using the posterior likelihood (see Supplementary materials).

The goodness of fit for each model was quantified using the integrated BIC (iBIC) (Guitart-Masip et al., 2012; Huys et al., 2011). Lower iBIC values indicate a better fit.

### Model recovery

We conducted model recovery for Effort task and RLWM task. The model recovery was done by generating 1000 synthetic datasets for each model by sampling mean and variance of the estimated parameter values observed for that model in our data. After that, we fitted all three models to the three generated synthetic datasets using the same hierarchical fitting as for the real data. For each synthetic subject in a given model, we quantified the goodness of the fit of each model using iBIC. See below for confusion matrices.

**For the effort task**, we tested 3 models, including the preregistered one (see Methods > *Computational models*). The model with a parabolic discounting term (constrained quadratic model) had the lowest bici (12868) compared to the preregistered model (13123) and our third candidate model (13004). We then calculated the winning model (the lowest BIC) for each synthetic dataset. We repeated this procedure for all the 1000 datasets and built a confusion matrix including the proportion of the synthetic datasets (simulated model) that were best fit by a given (fitted) model (see Figure S8).

**For the RLWM task**, we evaluated the alternative WMH model using the same procedure as for the models of the effort task models. While the WMH model yielded marginally lower bici (63085) than the original RLWM model (63419), we retained the originally hypothesized RLWM model. This decision was based on the parameter recovery which was superior to the RLWM model compared to the WMH model and the confusion matrix which showed that these models are both recoverable (see Figure S9).

### Two-step fitting procedure of the RLWM task’s model

Due to the discrete nature of the WM capacity parameter K in the RLWM task, we used a two-step fitting procedure.

In the first step, we estimated the parameter K for each participant by estimating all model parameters with a maximum likelihood (ML) approach separately for each of the possible values of K (1, 2, 3, 4 and 5). For each value of K, the model was estimated 10 times with different starting values. During this first step, the inverse temperature parameter β was fixed at 100 instead of 10. After estimating the parameters for every value of K, the K resulting in the smallest PL was chosen for a given participant.

In the second fitting step, the model was fitted hierarchically to the data as described above, using the K parameters obtained for each participant in the first step.
