## Supplementary figures and images for "Assessing reinforcement learning and vigor with an online decision-making battery"

### Figure S1

**Figure S1**

*Histograms of the parameters of interest for Session 1*

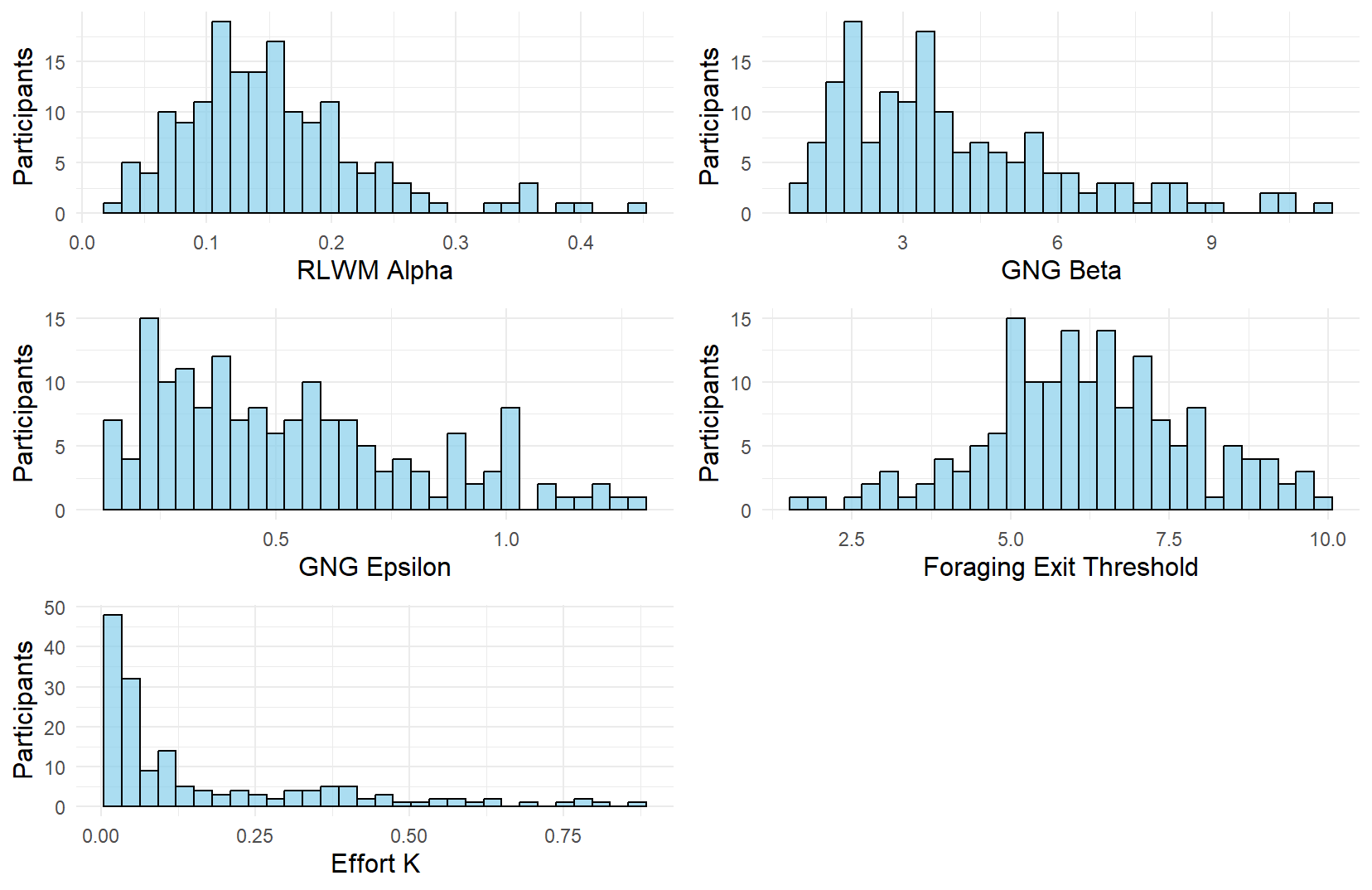

### Figure S3

**Figure S3**

*Histograms of all questionnaires for Session 1*

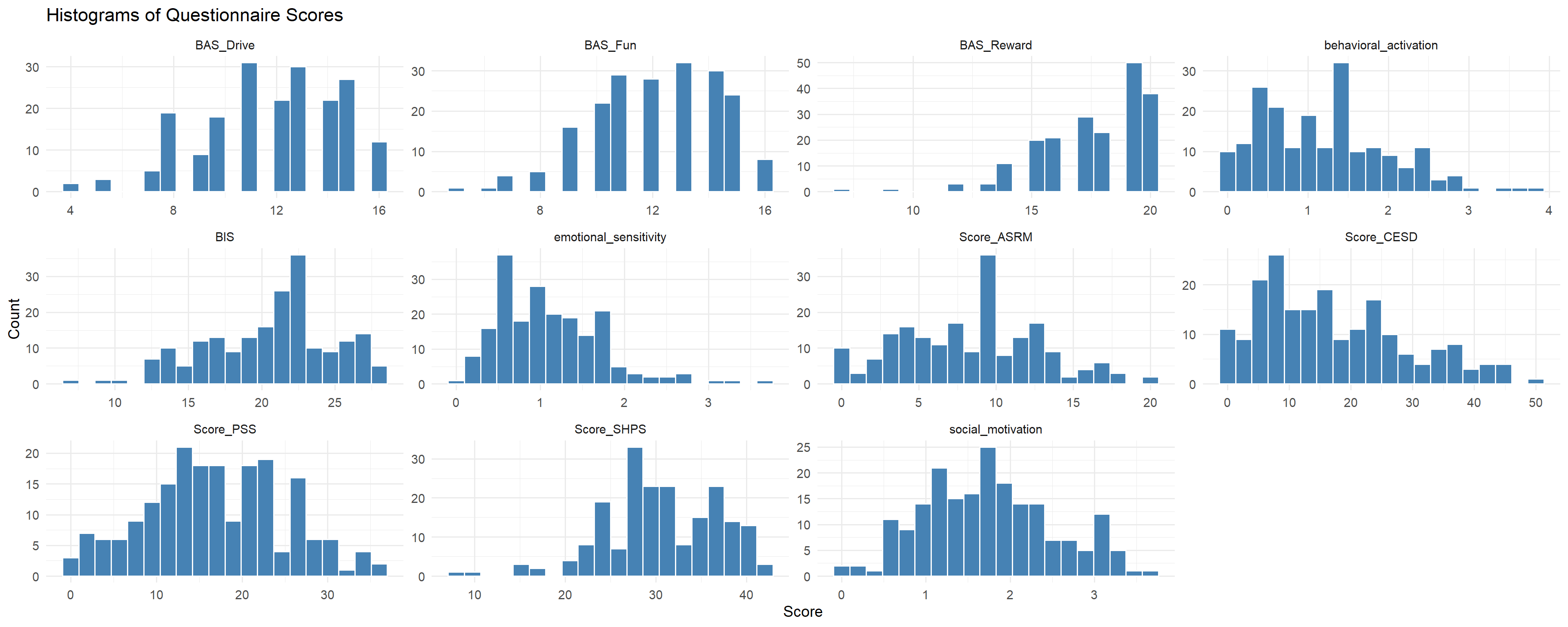

### Figure S4

**Figure S4**

*Parameter recovery for RLWM Task*

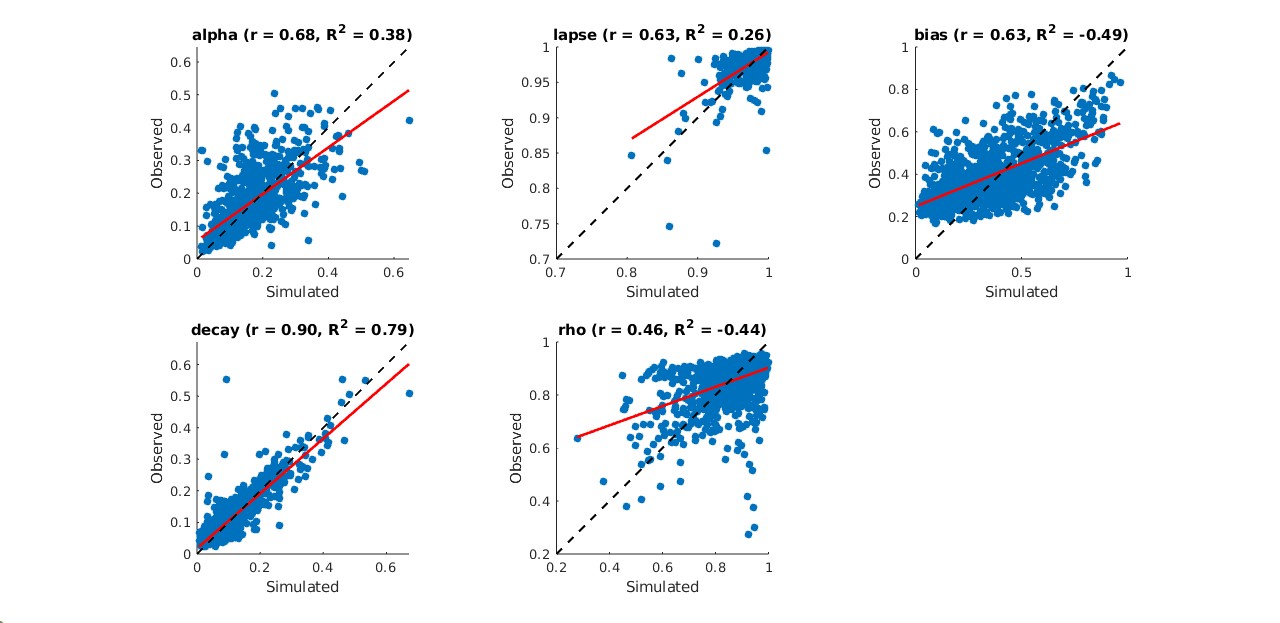

### Figure S5

**Figure S5**

*Parameter recovery for RLWM Task using WMH model*

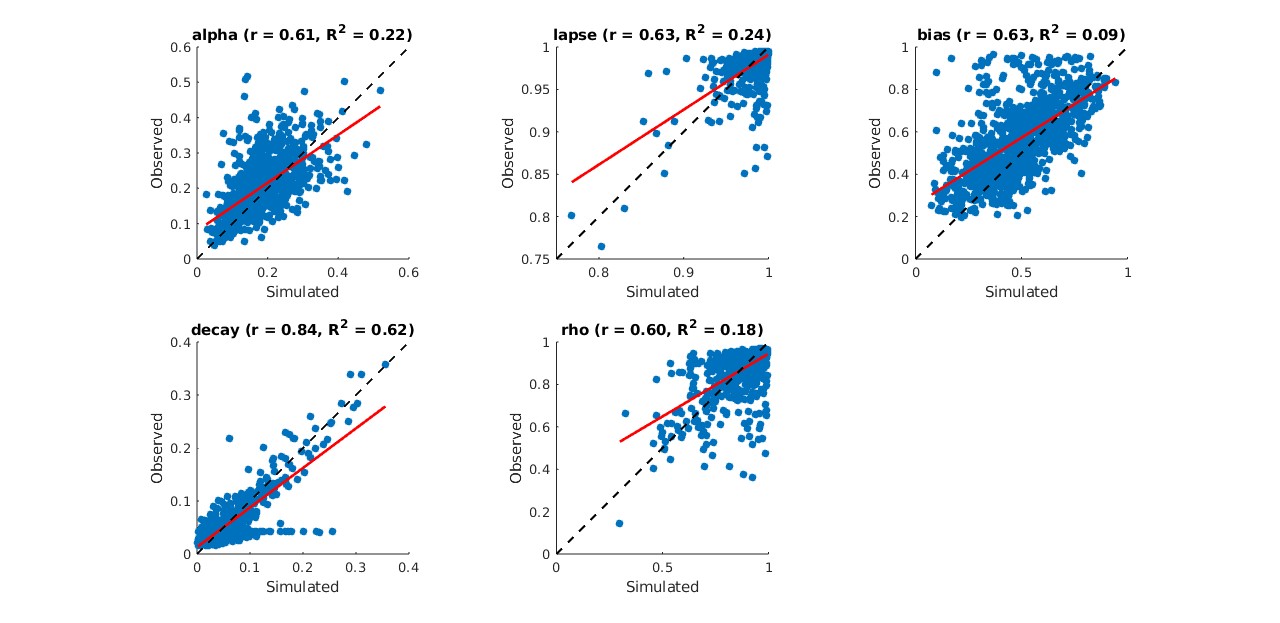

### Figure S6

**Figure S6**

*Parameter recovery for Effort Task*

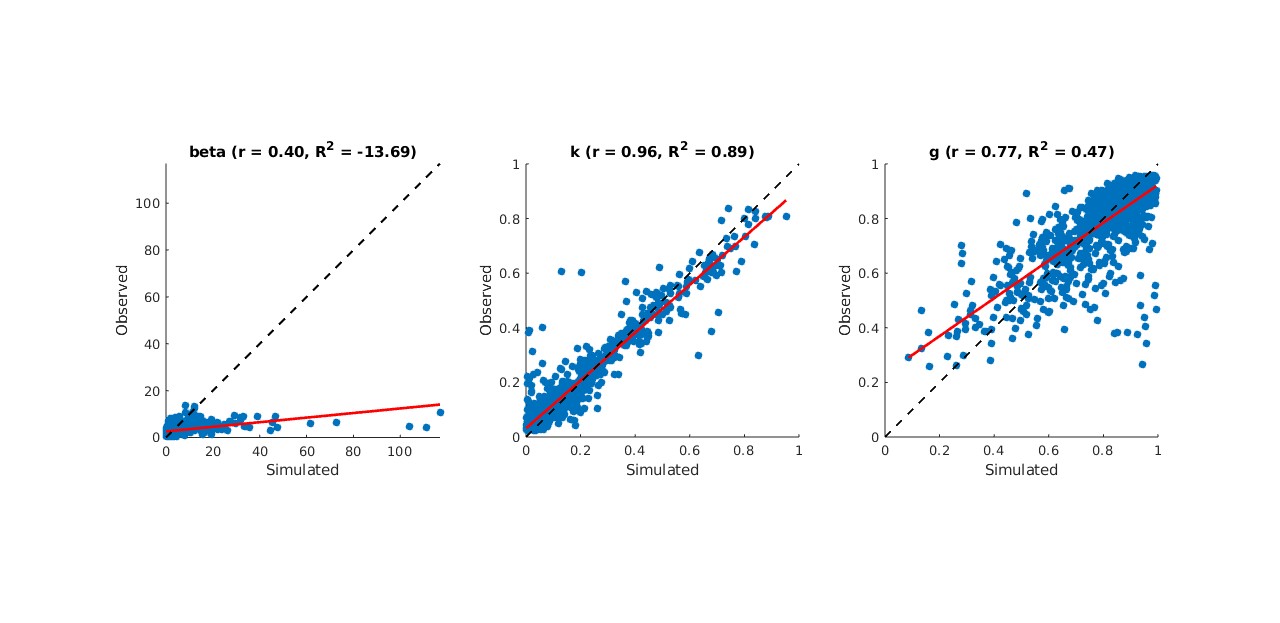

### Figure S7

**Figure S7**

*Parameter recovery for GNG Task*

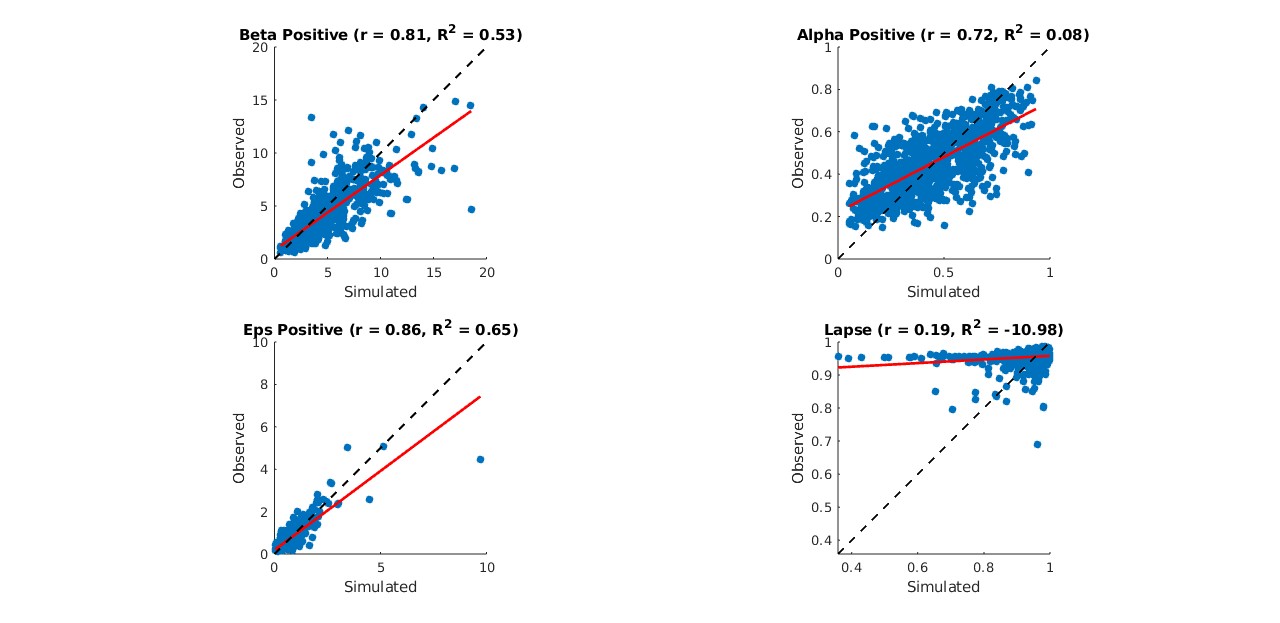

### Figure S8

**Figure S8**

*Confusion matrix for the models of the effort task*

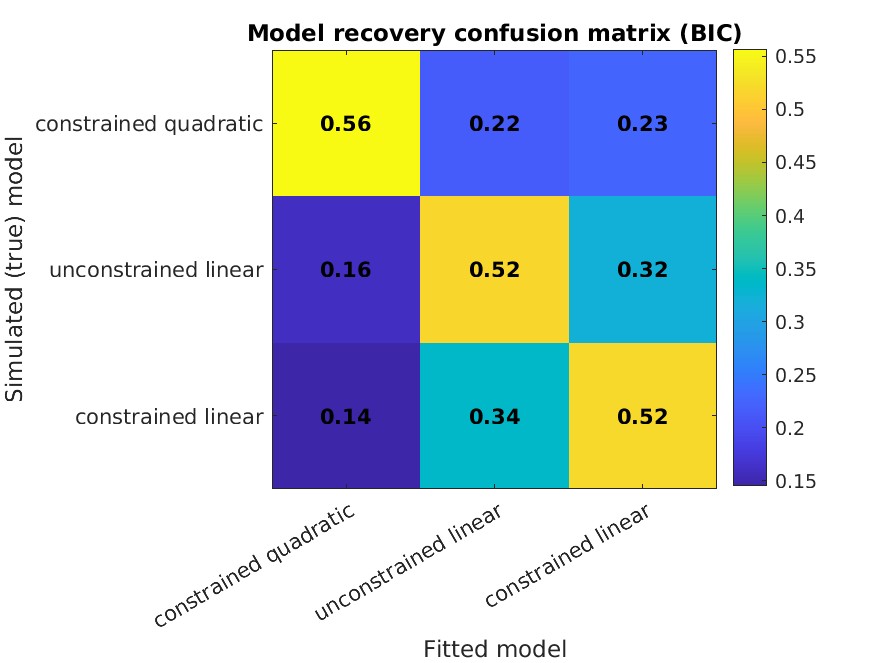

### Figure S9

**Figure S9**

*Confusion matrix for the WMH and RLWM models of the RLWM task*

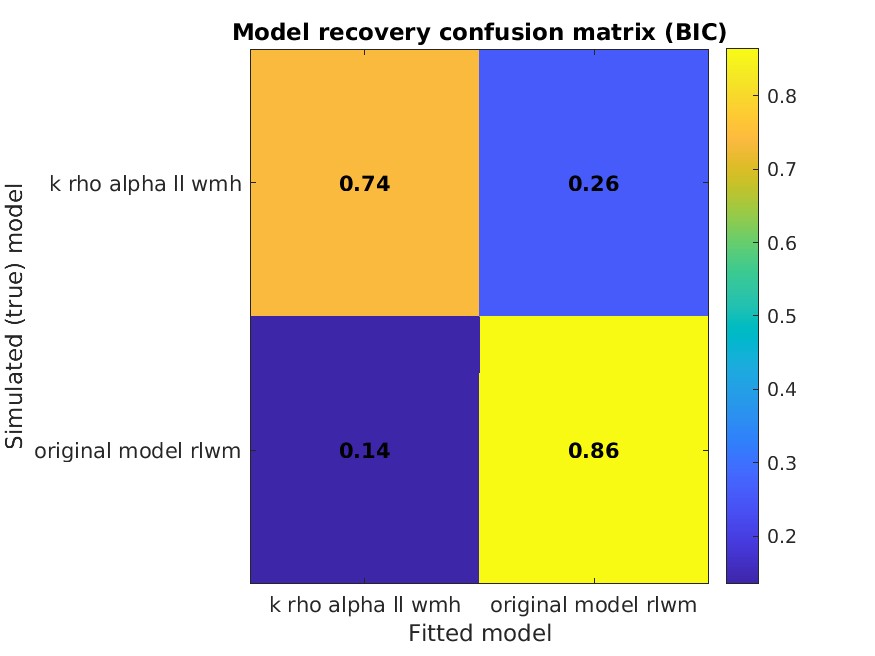
