## Supplementary material for "Assessing reinforcement learning and vigor with an online decision-making battery": Figure S10

*Preregistered outlier detection method using z-scores > |3| identified 6 outlier points out of which 5 were unique.*

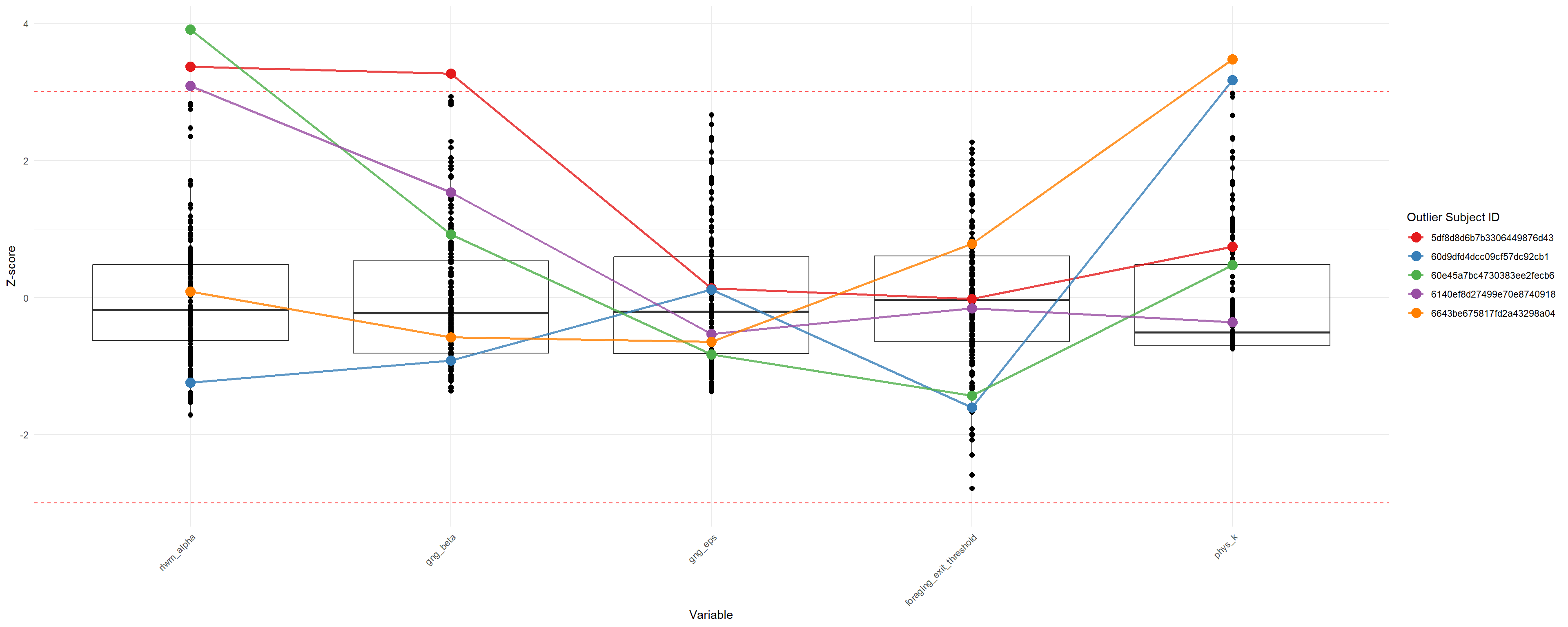
