## Supplementary material for "Assessing reinforcement learning and vigor with an online decision-making battery": Figure S11

*Outlier detection using the robust skewness-adjusted method identified 10 outlier points out of which 10 were unique.*

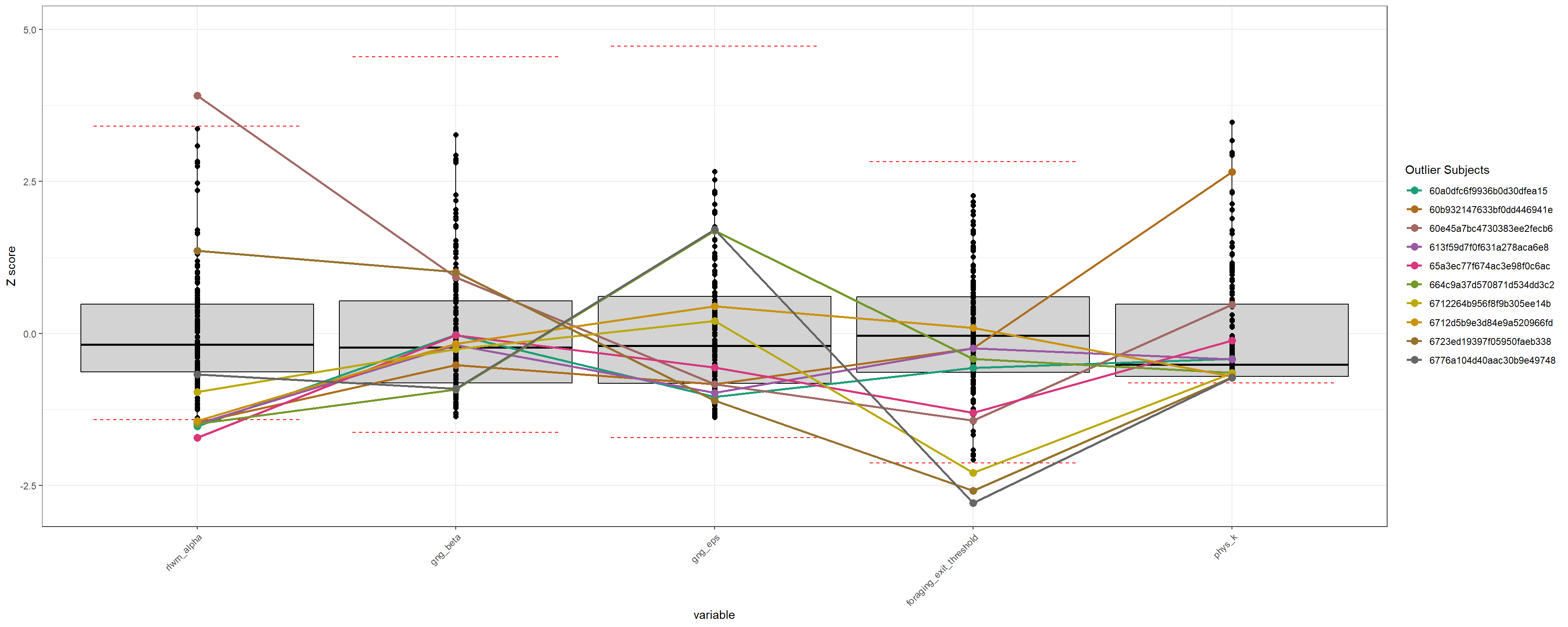
