## Supplementary material for "Assessing reinforcement learning and vigor with an online decision-making battery": Figure S12

*Multivariate outlier detection using Mahalanobis distance identified 9 outlier participants.*

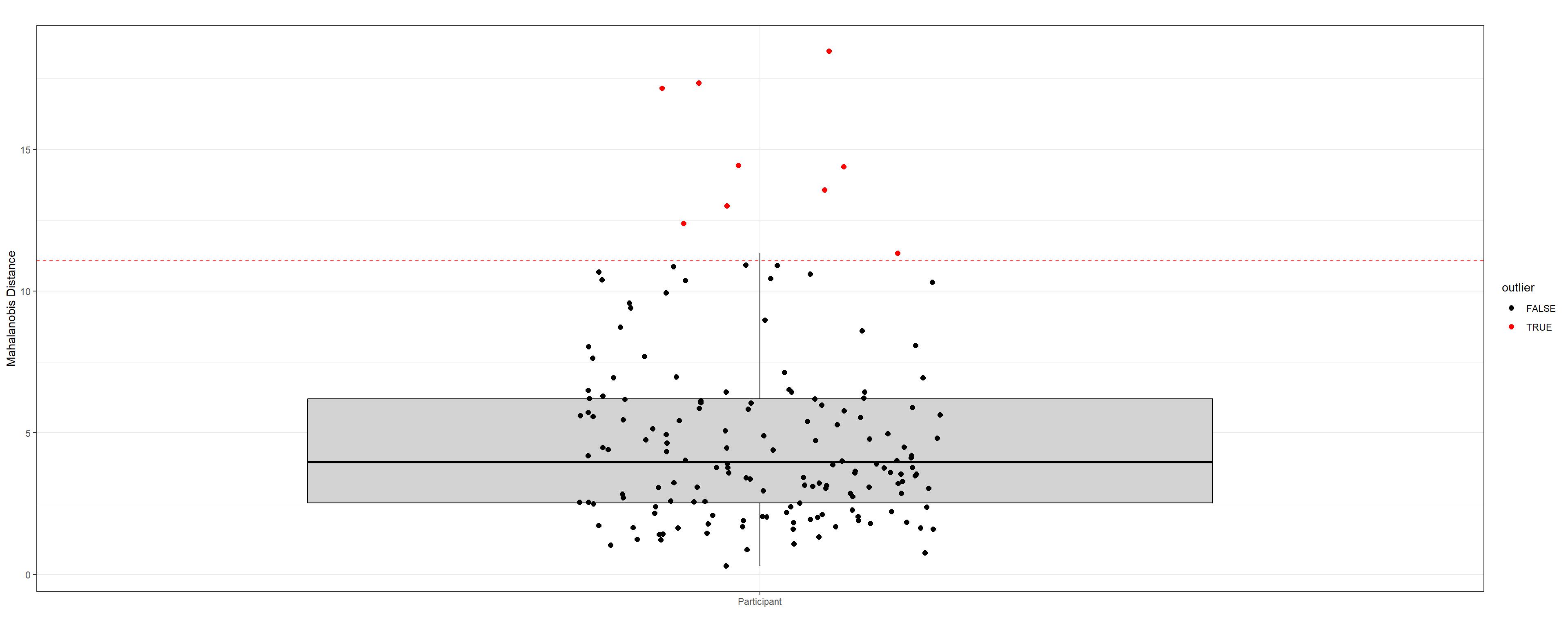
