## Supplementary material for "Assessing reinforcement learning and vigor with an online decision-making battery": Figure S14

*Spearman correlation matrix of computational parameters and questionnaire scores*

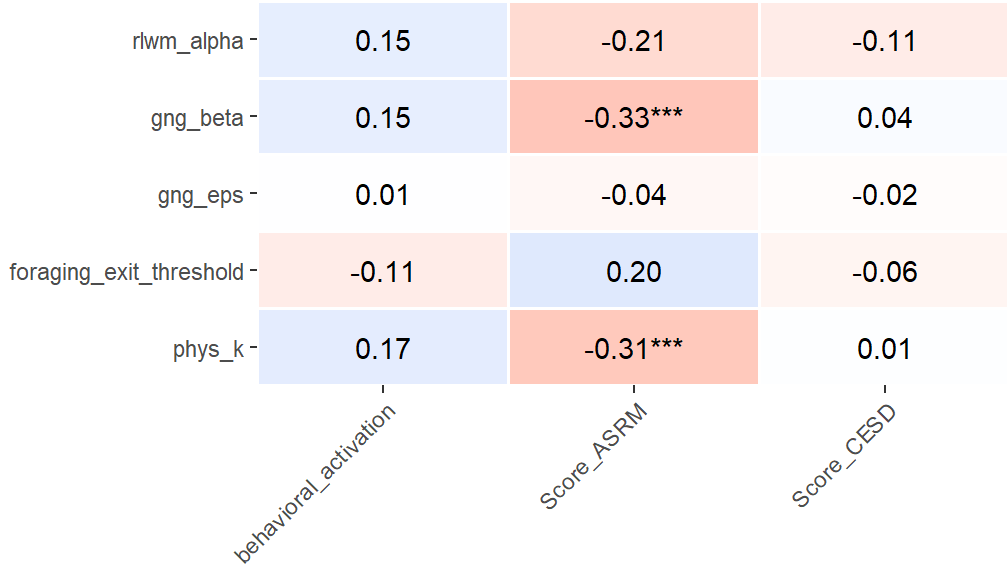

*Note.* phys_k = Effort κ; Applied Holm correction for multiple comparisons. *** p < .001; ** p < .01; * p < .05.
