## Supplementary material for "Assessing reinforcement learning and vigor with an online decision-making battery": Figure S15

*Spearman correlation matrix of RL factor and AMI items*

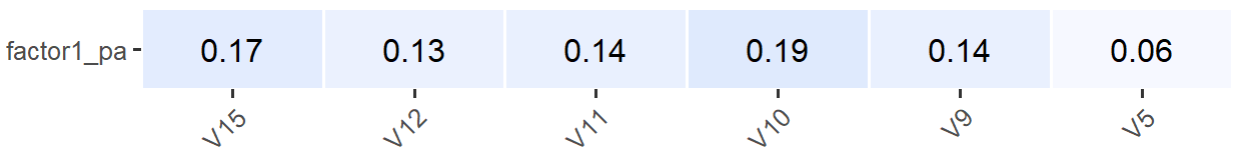

*Note.* Applied Holm correction for multiple comparisons. *** p < .001; ** p < .01; * p < .05.
