## Supplementary material for "Assessing reinforcement learning and vigor with an online decision-making battery": LPA

### BCH analysis of ES and SM subscales of AMI

**Emotional sensitivity**

For Emotional sensitivity, there were no statistically significant differences between the three profiles (p > .176).

Profile 1 (M = -0.15; SE = 0.14) and Profile 2 (M = 0.14; SE = 0.16) showed symmetrical values compared to the sample mean. Profile 3 (M = 0.04; SE = 0.12) had a mean value very close to the sample mean.

**Social motivation**

There was a significant difference in the sample means of Social motivation, only between Profile 1 and 3 (p = .008). There was no significant difference between profiles 2 and 3 (p = .051) and between Profile 1 and Profile 2 (p = .550).

Profile 3 exhibited the highest levels of social motivation (M = 0.30, SE = 0.13), whereas Profile 1 exhibited the lowest one (M = -0.19, SE = 0.12), while Profile 2 exhibited a value slightly below the sample mean (M = -0.08, SE = 0.14).
