## Supplementary material for "Assessing reinforcement learning and vigor with an online decision-making battery": Outliers

To investigate whether and how the EFA structure was sensitive to removal of outliers, we quantified univariate and multivariate outliers and re-ran the EFA after removing them.

### Univariate outliers

To quantify univariate outliers, we deviated from the preregistered method of normalizing the parameters to z-scores and labelling all values |z| > 3 as outliers. This is because the z-score method assumes that the variables fulfill the assumption of univariate normality, which is not a condition for EFA (Fabrigar et al., 1999; M. W. Watkins, 2018). Some of our parameters were not normally distributed and skewed (see Figure S1 in Supplementary Materials) and under these conditions many regular observations may be falsely flagged as outliers. However, for completeness, see Table S8 in Supplementary Materials. To circumvent this problem, it has been suggested that outliers should be defined based on interquartile range adjusted for skewed distribution (Hubert & Vandervieren, 2008). Excluding 10 outliers using this robust method preserved the original factor structure (see Table S9 in Supplementary Materials).

### Multivariate outliers

To identify multivariate outliers, we used Mahalanobis distance. Using the preregistered significance threshold of p < .05, based on the chi-square distribution with degrees of freedom equal to the number of variables, we identified 9 outliers. Their removal did not affect the EFA structure (see Table S10 in Supplementary Materials).
