## Supplementary material for "Assessing reinforcement learning and vigor with an online decision-making battery": Questionnaires

The other three other questionnaires that were included in our preregistration but not included in our main analyses were: Snaith-Hamilton Pleasure Scale (SHAPS), Behavioral Inhibition/Behavioral Activation Scales (BIS/BAS) and The Perceived Stress Scale (PSS).

### SHAPS

This questionnaire quantifies the hedonic capacity or, in other words, an ability to perceive pleasure of an individual through a self-assessment scale. It was developed to address needs for a simple scale independent of sociodemographic variables (Snaith et al., 1995). The 14-item questionnaire covered several domains of pleasure: social interaction, food and drink, sensory experiences, achievement and pastimes. Participants ranked each situation on a four-point scale to which degree did it caused them pleasure in the last few days. An example statement is: “I would be able to enjoy a beautiful landscape or view”.

### BIS/BAS

The questionnaire assesses the two personality systems. First, the aversive motivational system: behavioral inhibition system (BIS) which inhibits movement toward goals. Second, the appetitive motivational system: behavioral approach system (BAS) that should reflect the proneness to engage in goal-directed efforts (Carver & White, 1994). The scale contains 24 statements that are rated by participants based on whether they agree or disagree with. The scale is then divided into BIS items and BAS items with specific subscales Drive, Fun Seeking and Reward Responsiveness.

### PSS

This scale was developed in order to measure the degree to which situations in one’s life are appraised as stressful (Cohen et al., 1983). The questions ask about feelings and thoughts during the last month. Participants answer questions based on given frequency: how they felt or thought from “Never” to “Very often”. The scale consists of 10 questions such as “In the last month, how often have you felt that things were going your way?” or “In the last month, how often have you felt that you were on top of things?”.
