## Supplementary material for "Assessing reinforcement learning and vigor with an online decision-making battery": Regression analyses

To investigate the associations of the factors and the questionnaire scores, adjusting for *age*, *sex*, and the other *factor*, we conducted a series of exploratory multiple regressions, one for each questionnaire score. We found that *factor 1* (RL) significantly predicted the *BA* score, *t*(156) = 2.46, β = 0.22, *p* = .015) when controlling for age and sex. This was a surprising finding given the non-significant correlation between BA and RL (ρ = .19; *p* = .116). Further, the *ASRM* score was predicted by *age*, *t*(156) = -3.24, β = -0.10, *p* = .001 and *factor 1* (RL), *t*(156) = -4.17, β = -1.90, *p* < .001, but not by *factor 2* *t*(156) = 1.63, β = 1.13, *p* = .105. To further explore what influenced the *factor 2* not being associated with the *ASRM* score (despite the significant univariate correlation reported above), we ran another regression with *age* and *sex* but without *factor 1*. In this regression, *factor 2* significantly predicted the ASRM score, *t*(157) = 2.32, β = 1.66, *p* = .022. This implied some shared variance between RL and vigor in which RL may dominate when included in the analysis together with vigor. The *CESD* scores were not predicted by any of the demographic variables (*p* > .132) or factor scores (*p* > .912).
