## Supplementary material for "Assessing reinforcement learning and vigor with an online decision-making battery": Sample

### Demographics

Participants in Session 1 came originally from 23 countries, majority of them from South Africa (24.07%), the United Kingdom (24.07%), and the United States (11.73%). Most of the participants were employed: full-time (56.17%) and part-time (12.35%). 11.11% of participants were unemployed, and 20.99% of participants had some other arrangement*.* 53.70% of participants reported being students. The age ranged from 19 to 60 (M = 32.99, SD = 10.23). The range of the number of previously finished and approved submissions in other studies on Prolific ranged from 13 to 8728, with a median of 417.50.

In Session 2, participants came originally from 18 countries; majority of participants from the United Kingdom (35.21%), followed by South Africa (21.13%). Majority of the sample were employed full-time (61.97%) and 30.99% were reported students. The age range of the sample spanned from 19 to 58 (M = 33.56, SD = 10.68). The range of the number of previously finished and approved submissions in other studies on Prolific ranged from 16 to 8728, with a median of 454.00.

### Sample size rationale

Based on our pilot data, we carried out a power analysis for the predictive validity checks. These are based on correlations between the factor scores from the EFA (Vigor and RL) and the questionnaire data. In our pilot sample (*N* = 53), we observed an average correlation of approximately r = ±0.30 between vigor and BA (as measured with the AMI questionnaire) and vigor and mania symptoms (as measuread by ASRM). Based on the power analysis done using an R package *pwr* (Champely, 2020), we needed at least 85 participants to detect a correlation with an *r* = 0.3 with *α* = 0.05 (two-sided) and power = 0.80. To increase our power to detect associations with external factors, account for potential exclusions, and allow for robust comparisons (including test-retest reliability), we recruited 200 participants for Session 1 and 100 participants for Session 2. In fact, with 200 participants, we were able to detect a Pearson correlation of *r* = 0.20, with *α* = 0.05 (two-sided), and power = 0.80.
