## Supplementary material for "Assessing reinforcement learning and vigor with an online decision-making battery": Table S1

*Reported country of residence in the analyzed sample*

| **Country of Residence** | **Session 1 (N = 162)** | **Session 2 (N = 71)** |
| --- | --- | --- |
| Algeria | 0.62% | 1.41% |
| Australia | 0.62% | 0.00% |
| Belgium | 0.62% | 1.41% |
| Brazil | 1.85% | 2.82% |
| Canada | 2.47% | 2.82% |
| Chile | 1.85% | 0.00% |
| Denmark | 0.62% | 0.00% |
| France | 1.23% | 0.00% |
| Germany | 3.09% | 1.41% |
| Greece | 3.70% | 2.82% |
| Hungary | 3.70% | 5.63% |
| Ireland | 0.62% | 1.41% |
| Italy | 4.32% | 5.63% |
| Kenya | 1.85% | 2.82% |
| Mexico | 1.85% | 1.41% |
| Netherlands | 0.62% | 0.00% |
| Poland | 4.32% | 5.63% |
| Portugal | 3.09% | 0.00% |
| Slovenia | 0.62% | 0.00% |
| South Africa | 24.07% | 21.13% |
| Spain | 2.47% | 1.41% |
| United Kingdom | 24.07% | 35.21% |
| United States | 11.73% | 7.04% |
