## Supplementary material for "Assessing reinforcement learning and vigor with an online decision-making battery": Table S2

*Models that entered LPA analysis*

| **Name** | **Profiles** | **Par.** | **AIC** | **BIC** | **Entropy** | **P. (min, max)** | **N (min/max)** |
| --- | --- | --- | --- | --- | --- | --- | --- |
| equal var 1 | 1 | 10 | 2313.67 | 2344.54 | 1.00 | [1.00, 1.00] | [1.00, 1.00] |
| equal var 2 | 2 | 16 | 2243.24 | 2292.64 | 0.93 | [0.97, 0.98] | [0.23, 0.77] |
| equal var 3 | 3 | 22 | 2246.66 | 2314.59 | 0.81 | [0.86, 0.94] | [0.15, 0.65] |
| equal var 4 | 4 | 28 | 2181.45 | 2267.90 | 0.89 | [0.82, 0.98] | [0.04, 0.62] |
| equal var 5 | 5 | 34 | 2179.10 | 2284.08 | 0.80 | [0.76, 0.97] | [0.04, 0.41] |
| equal var 6 | 6 | 40 | 2176.84 | 2300.35 | 0.82 | [0.70, 0.99] | [0.04, 0.43] |
| free var 1 | 1 | 10 | 2313.67 | 2344.54 | 1.00 | [1.00, 1.00] | [1.00, 1.00] |
| free var 2 | 2 | 21 | 2227.20 | 2292.04 | 0.61 | [0.78, 0.95] | [0.35, 0.65] |
| ***free var 3** | **3** | **32** | **2001.19** | **2099.99** | **0.78** | **[0.85, 0.96]** | **[0.31, 0.38]** |
| Equal var equal cov 1 | 1 | 20 | 2301.01 | 2362.76 | 1.00 | [1.00, 1.00] | [1.00, 1.00] |
| equal var equal cov 2 | 2 | 26 | 2225.91 | 2306.19 | 0.96 | [0.98, 1.00] | [0.23, 0.77] |
| equal var equal cov 3 | 3 | 32 | 2256.49 | 2355.29 | 0.81 | [0.87, 0.94] | [0.18, 0.62] |
| equal var equal cov 4 | 4 | 38 | 2185.20 | 2302.52 | 0.90 | [0.88, 0.97] | [0.05, 0.60] |
| equal var equal cov 5 | 5 | 44 | 2210.58 | 2346.43 | 0.86 | [0.90, 0.93] | [0.08, 0.35] |
| equal var equal cov 6 | 6 | 50 | 2184.07 | 2338.45 | 0.82 | [0.70, 0.99] | [0.04, 0.30] |
| free var equal cov 1 | 1 | 20 | 2301.01 | 2362.76 | 1.00 | [1.00, 1.00] | [1.00, 1.00] |
| free var equal cov 2 | 2 | 31 | 2038.74 | 2134.45 | 0.87 | [0.95, 0.98] | [0.45, 0.55] |
| free var equal cov 3 | 3 | 42 | 2010.75 | 2140.43 | 0.80 | [0.87, 0.99] | [0.30, 0.40] |
| free var equal cov 4 | 4 | 53 | 1957.79 | 2121.43 | 0.84 | [0.79, 0.99] | [0.15, 0.38] |
| free var equal cov 5 | 5 | 64 | 1963.37 | 2160.97 | 0.86 | [0.72, 1.00] | [0.03, 0.38] |
| free var equal cov 6 | 6 | 75 | 1968.97 | 2200.54 | 0.87 | [0.75, 1.00] | [0.04, 0.33] |
| equal var free cov 1 | 1 | 20 | 2301.01 | 2362.76 | 1.00 | [1.00, 1.00] | [1.00, 1.00] |
| equal var free cov 2 | 2 | 36 | 2239.40 | 2350.56 | 0.95 | [0.98, 0.99] | [0.23, 0.77] |
| equal var free cov 3 | 3 | 52 | 2222.92 | 2383.48 | 0.92 | [0.92, 0.97] | [0.10, 0.73] |
| equal var free cov 4 | 4 | 68 | 2207.47 | 2417.43 | 0.91 | [0.94, 0.97] | [0.10, 0.61] |
| equal var free cov 5 | 5 | 84 | 2227.35 | 2486.71 | 0.84 | [0.87, 0.95] | [0.05, 0.46] |
| equal var free cov 6 | 6 | 100 | 2201.14 | 2509.90 | 0.90 | [0.91, 1.00] | [0.04, 0.30] |
| free var  free cov 1 | 1 | 20 | 2301.01 | 2362.76 | 1.00 | [1.00, 1.00] | [1.00, 1.00] |
| free var  free cov 2 | 2 | 41 | 2229.50 | 2356.09 | 0.71 | [0.89, 0.92] | [0.42, 0.58] |
| free var  free cov 3 | 3 | 62 | 2050.10 | 2241.53 | 0.86 | [0.92, 0.99] | [0.15, 0.44] |
| free var  free cov 4 | 4 | 83 | 1982.53 | 2238.81 | 0.91 | [0.92, 1.00] | [0.07, 0.41] |
| free var free cov 5 | 5 | 104 | 2025.73 | 2346.84 | 0.92 | [0.92, 1.00] | [0.05, 0.41] |
| free var free cov 6 | 6 | 125 | 2010.26 | 2396.21 | 0.93 | [0.91, 1.00] | [0.06, 0.33] |

*Note.* Name = Model specifications (e.g. equal vs. free variances); Profiles = Number of latent profiles estimated in the model; Par. **=** Number of free parameters estimated in the model; AIC = Akaike Information Criterion; BIC = Bayesian Information Criterion; Entropy = values closer to 1.00 indicate better separation between profiles; P. (min, max) = Minimum and maximum average posterior assignment probabilities across profiles; n (min, max) = Proportion of the total sample assigned to the smallest and largest profiles respectively; * = Best model based on the lowest BIC**:** During the fitting procedure, we received a warning regarding the FreeVar3 model’s potential lack of stability. To rule out the possibility of converging at a local minimum and to assess the model overall convergence and stability, we repeatedly re-estimated the same model using 50 different random starting values using the function *mxTryHard().* The resulting log-likelihoods were identical across all re-estimated models, indicating that the initial solution corresponded to a global minimum. This analysis demonstrated reliability and stability of the FreeVar3 model; *All models with n_min_ < 0.1 were not included in further LPA analyses.*
