## Supplementary material for "Assessing reinforcement learning and vigor with an online decision-making battery": Table S3

*Parameter estimates and significance levels for the FreeVar3 Latent Profile Analysis model.*

| Label | Estimate | SE | *p* | 95% CI | Profile |
| --- | --- | --- | --- | --- | --- |
| Means.rlwm_alpha | -0.24* | 0.12 | 0.04 | [-0.47, -0.01] | profile1 |
| Means.gng_beta | -0.42*** | 0.09 | 0.00 | [-0.60, -0.23] | profile1 |
| Means.gng_eps | 0.28 | 0.15 | 0.06 | [-0.01, 0.58] | profile1 |
| Means.foraging_exit_threshold | 0.17 | 0.16 | 0.29 | [-0.14, 0.48] | profile1 |
| Means.effort_k | -0.70*** | 0.01 | 0.00 | [-0.71, -0.69] | profile1 |
| Variances.rlwm_alpha | 0.67*** | 0.15 | 0.00 | [0.38, 0.96] | profile1 |
| Variances.gng_beta | 0.33*** | 0.10 | 0.00 | [0.14, 0.53] | profile1 |
| Variances.gng_eps | 1.25*** | 0.23 | 0.00 | [0.80, 1.69] | profile1 |
| Variances.foraging_exit_threshold | 1.50*** | 0.29 | 0.00 | [0.94, 2.06] | profile1 |
| Variances.effort_k | 0.00** | 0.00 | 0.01 | [0.00, 0.00] | profile1 |
| Means.rlwm_alpha | -0.29** | 0.11 | 0.01 | [-0.50, -0.08] | profile2 |
| Means.gng_beta | -0.45*** | 0.10 | 0.00 | [-0.64, -0.26] | profile2 |
| Means.gng_eps | 0.14 | 0.17 | 0.42 | [-0.20, 0.48] | profile2 |
| Means.foraging_exit_threshold | -0.19 | 0.14 | 0.17 | [-0.46, 0.08] | profile2 |
| Means.effort_k | 0.80*** | 0.20 | 0.00 | [0.41, 1.19] | profile2 |
| Variances.rlwm_alpha | 0.37*** | 0.10 | 0.00 | [0.18, 0.56] | profile2 |
| Variances.gng_beta | 0.31*** | 0.08 | 0.00 | [0.16, 0.45] | profile2 |
| Variances.gng_eps | 0.98*** | 0.22 | 0.00 | [0.56, 1.41] | profile2 |
| Variances.foraging_exit_threshold | 0.74*** | 0.16 | 0.00 | [0.42, 1.06] | profile2 |
| Variances.effort_k | 1.29*** | 0.27 | 0.00 | [0.76, 1.82] | profile2 |
| Means.rlwm_alpha | 0.56** | 0.20 | 0.01 | [0.17, 0.94] | profile3 |
| Means.gng_beta | 0.91*** | 0.21 | 0.00 | [0.49, 1.33] | profile3 |
| Means.gng_eps | -0.46*** | 0.10 | 0.00 | [-0.66, -0.27] | profile3 |
| Means.foraging_exit_threshold | -0.02 | 0.12 | 0.84 | [-0.27, 0.22] | profile3 |
| Means.effort_k | 0.08 | 0.13 | 0.56 | [-0.19, 0.34] | profile3 |
| Variances.rlwm_alpha | 1.50*** | 0.32 | 0.00 | [0.89, 2.12] | profile3 |
| Variances.gng_beta | 1.20*** | 0.27 | 0.00 | [0.68, 1.73] | profile3 |
| Variances.gng_eps | 0.38*** | 0.09 | 0.00 | [0.21, 0.55] | profile3 |
| Variances.foraging_exit_threshold | 0.56*** | 0.14 | 0.00 | [0.29, 0.84] | profile3 |
| Variances.effort_k | 0.71*** | 0.16 | 0.00 | [0.40, 1.01] | profile3 |
| mix3.weights[1,1] | 1.00 |  |  |  |  |
| mix3.weights[1,2] | 0.79*** | 0.19 | 0.00 | [0.42, 1.17] |  |
| mix3.weights[1,3] | 0.85*** | 0.22 | 0.00 | [0.42, 1.29] |  |

*Note.* The specific model parameter being estimated (e.g., means of decision-making variables); Estimate **=** estimated value of the parameter. Asterisks indicate the level of statistical significance: *** p < .001; ** p < .010; * p < .050; SE = Standard Error of the estimate; p = p-value; 95% CI = 95% Confidence Interval for the estimate. If the interval does not include zero, the effect is considered significant; Profile = The specific latent profile to which the estimates refer (e.g., Profile 1).
