## Supplementary material for "Assessing reinforcement learning and vigor with an online decision-making battery": Table S4

| *Descriptive Statistics of All Parameters* | | | | | | | | |
| --- | --- | --- | --- | --- | --- | --- | --- | --- |
| Parameter | M | SD | SE | Median | Q5 | Q25 | Q75 | Q95 |
| Foraging Exit Threshold | 6.22 | 1.63 | 0.13 | 6.17 | 3.50 | 5.18 | 7.21 | 8.99 |
| RLWM α | 0.15 | 0.07 | 0.01 | 0.14 | 0.06 | 0.11 | 0.19 | 0.28 |
| RLWM Bias | 0.37 | 0.16 | 0.01 | 0.33 | 0.17 | 0.23 | 0.48 | 0.68 |
| RLWM Decay | 0.09 | 0.05 | 0.00 | 0.09 | 0.03 | 0.05 | 0.13 | 0.19 |
| RLWM Lapse | 0.99 | 0.01 | 0.00 | 0.99 | 0.96 | 0.98 | 0.99 | 0.99 |
| RLWM ρ | 0.88 | 0.08 | 0.01 | 0.91 | 0.68 | 0.86 | 0.93 | 0.94 |
| GNG α | 0.44 | 0.16 | 0.01 | 0.44 | 0.19 | 0.32 | 0.58 | 0.70 |
| GNG Β | 3.96 | 2.20 | 0.17 | 3.46 | 1.44 | 2.17 | 5.13 | 8.32 |
| GNG Epsilon | 0.53 | 0.28 | 0.02 | 0.48 | 0.18 | 0.30 | 0.70 | 1.03 |
| GNG Lapse | 0.96 | 0.03 | 0.00 | 0.96 | 0.93 | 0.96 | 0.97 | 0.98 |
| Effort Β | 2.81 | 2.09 | 0.16 | 2.18 | 0.40 | 0.97 | 4.63 | 6.16 |
| Effort K | 0.17 | 0.20 | 0.02 | 0.07 | 0.02 | 0.03 | 0.27 | 0.60 |
| Effort Lapse | 0.81 | 0.16 | 0.01 | 0.86 | 0.40 | 0.76 | 0.92 | 0.95 |
