## Supplementary material for "Assessing reinforcement learning and vigor with an online decision-making battery": Table S5

| **Table S5**  *Factor Loadings for Session 1* | | |
| --- | --- | --- |
| Parameter | RL | Vigor |
| RLWM α | .54 |  |
| GNG β | .58 |  |
| GNG ε | -.36 | .20 |
| Foraging Exit Threshold |  | .35 |
| Effort κ | .11 | -.32 |
