## Supplementary material for "Assessing reinforcement learning and vigor with an online decision-making battery": Table S6

| **Table S6**  *EFA when using the preregistered ML extraction method* | | |
| --- | --- | --- |
| Parameter | RL | Vigor |
| RLWM α | .53 |  |
| GNG β | .59 |  |
| GNG ε | -.36 | .20 |
| Foraging Exit Threshold |  | .36 |
| Effort κ | .11 | -.32 |

*Note.* Variance explained 15% for factor 1 and 5% for factor 2.
