## Supplementary material for "Assessing reinforcement learning and vigor with an online decision-making battery": Table S7

*EFA of Session 2*

| Parameter | Factor 1 | Factor 2 |
| --- | --- | --- |
| RLWM α | .45 |  |
| GNG β | .75 | -.14 |
| GNG ε | -.15 | .28 |
| Foraging Exit Threshold | .12 | .40 |
| Effort κ |  | .19 |

*Note.* Variance explained 16% for factor 1 and 6% for factor 2.
