## Supplementary material for "Assessing reinforcement learning and vigor with an online decision-making battery": Table S8

*EFA after removing outliers using z-scores method*

| Parameter | RL | Vigor |
| --- | --- | --- |
| RLWM α | .46 |  |
| GNG β | .49 |  |
| GNG ε | -.43 |  |
| Foraging Exit Threshold |  | .55 |
| Effort κ | .20 | -.17 |

*Note.* Variance explained 14% for factor 1 and 7% for factor 2.
