## Supplementary material for "Assessing reinforcement learning and vigor with an online decision-making battery": Table S9

*EFA after removing outliers using the robust skewness-adjusted method*

| Parameter | RL | Vigor |
| --- | --- | --- |
| RLWM α | .55 |  |
| GNG β | .54 |  |
| GNG ε | -.35 | .16 |
| Foraging Exit Threshold |  | .52 |
| Effort κ | .12 | -.31 |

*Note.* Variance explained 15% for factor 1 and 8% for factor 2.
