## Supplementary material for "Assessing reinforcement learning and vigor with an online decision-making battery": Table S10

*EFA after removing multivariate outliers detected by the Mahalanobis distance*

| Parameter | RL | Vigor |
| --- | --- | --- |
| RLWM α | .48 |  |
| GNG β | .56 |  |
| GNG ε | -.39 | .26 |
| Foraging Exit Threshold |  | .36 |
| Effort κ |  | -.37 |

*Note.* Variance explained 14% for factor 1 and 7% for factor 2.
