## Supplementary material for "Assessing reinforcement learning and vigor with an online decision-making battery": Table S11

*EFA after substituting RLWM α with WMH α and WMH ρ*

| Parameter | RL | Vigor |
| --- | --- | --- |
| WMH α | .72 |  |
| WMH ρ | .58 |  |
| GNG β | .30 | -.25 |
| GNG ε | -.12 | .39 |
| Foraging Exit Threshold | .24 | .32 |
| Effort κ |  | -.28 |

*Note.* Variance explained 17% for factor 1 and 7% for factor 2.
