## Supplementary material for "Assessing reinforcement learning and vigor with an online decision-making battery": Table S12

*Factor analysis of Session 1 with participants that have done both Session 1 and Session 2 (N = 64).*

| Parameter | Factor 1 | Factor 2 |
| --- | --- | --- |
| RLWM α | .76 |  |
| GNG β | .34 | .32 |
| GNG ε | -.18 | .11 |
| Foraging Exit Threshold |  | .53 |
| Effort κ |  | .22 |

*Note.* Variance explained 15% for factor 1 and 8% for factor 2.
