## Supplementary material for "Assessing reinforcement learning and vigor with an online decision-making battery": Table S13

*Test-retest reliability of all individual parameters across sessions*

| Parameter | ICC | 95% CI |
| --- | --- | --- |
| **Foraging Exit Threshold** | **.76** | **[.63; .85]** |
| **RLWM α** | **.20** | **[-.03; .41]** |
| RLWM Bias | .35 | [.09**;** .56] |
| RLWM Decay | .59 | [.41**;** .73] |
| RLWM Lapse | .41 | [.19**;** .59] |
| RLWM ρ | .16 | [-.08**;** .38] |
| GNG **α** | .20 | [-.04**;** .42] |
| **GNG β** | **.73** | **[.59; .82]** |
| **GNG ε** | **.53** | **[.33; .68]** |
| GNG Lapse | .08 | [-.07**;** .25] |
| Effort β | .33 | [.09**;** .53] |
| **Effort κ** | **.64** | **[.47; .76]** |
| Effort Lapse | .01 | [-.13**;** .18] |

*Note.* ICC(2,1) for each parameter with the 95 % confidence interval (95% CI). Parameters in bold are included in the main analysis.
