## Supplementary material for "Assessing reinforcement learning and vigor with an online decision-making battery": Table S14

*3-factor solution for Session 1 of the pilot study including all parameters*

| Parameter | Inverse RL | Vigor | Biases |
| --- | --- | --- | --- |
| RLWM α | -.62 |  |  |
| RLWM Bias | .66 |  | .27 |
| RLWM Decay | .77 |  | .11 |
| RLWM Lapse |  |  | .21 |
| RLWM ρ | -.65 |  | .22 |
| GNG α+ | -.29 |  | .72 |
| GNG α- | -.31 | -.28 | .41 |
| GNG β+ | -.40 |  |  |
| GNG β- | .33 | .32 |  |
| GNG ε+ | .71 | .63 |  |
| GNG ε- | .42 | -.69 | -.11 |
| GNG Lapse | -.54 | .16 | .21 |
| Foraging Exit Threshold |  | .39 | -.11 |
| Effort β | -.19 | .16 | -.31 |
| Effort κ | .11 | -.53 | .29 |

*Note.* Variance explained 22% for factor 1, 10% for factor 2 and 8% for factor 3.
