## Supplementary material for "Assessing reinforcement learning and vigor with an online decision-making battery": Table S15

*2-factor solution for Session 1 of the pilot study including all parameters*

| Parameter | RL | Inverse vigor |
| --- | --- | --- |
| RLWM α | .63 | -.11 |
| RLWM Bias | .58 | .26 |
| RLWM Decay | -.71 | .31 |
| RLWM Lapse |  | .11 |
| RLWM ρ | .64 | -.18 |
| GNG α+ | .34 |  |
| GNG α- | .42 | .27 |
| GNG β+ | .35 | -.21 |
| GNG β- | -.39 | -.19 |
| GNG ε+ | -.36 | .19 |
| GNG ε- | -.79 | -.31 |
| GNG Lapse | .51 | -.23 |
| Foraging Exit Threshold | -.13 | -.40 |
| Effort β |  | -.29 |
| Effort κ |  | .62 |

*Note.* + indicates a parameter estimated from a reward condition, - indicates a parameter estimated from a punishment condition. Variance explained 21% for factor 1 and 11% for factor 2.
