## Supplementary material for "Assessing reinforcement learning and vigor with an online decision-making battery": Table S16

*2-factor solution for Session 1 of the pilot study including parameters of interest*

| Parameter | RL | Vigor |
| --- | --- | --- |
| RLWM α | .73 |  |
| GNG β | .69 |  |
| GNG ε | -.47 | .53 |
| Foraging Exit Threshold |  | .52 |
| Effort κ |  | -.49 |

*Note.* Variance explained 25% for factor 1 and 16% for factor 2.
