## Supplementary material for "Assessing reinforcement learning and vigor with an online decision-making battery": Tasks

### RLWM task

Participants were instructed that each correct key, associated with a particular stimulus, was independent from the other stimuli and that learning one response did not provide information for the other (i.e., a key “w” may be correct both for a “chair” and an “elephant” or may not be correct for any stimuli within a stimulus set).

Participants earned a point for each correct response and did not collect any points for an incorrect one. The total number of collected points was converted into a performance bonus at the end of the task.

The task consisted of 480 trials divided into 2 blocks (240 trials each). Each of the blocks was made up of mini-blocks. Each mini-block varied in the number of stimuli for which participants needed to learn a response (set size), manipulating WM load. These different WM loads helped to differentiate the contributions of RL and WM to task performance. In total, there were 5 levels of WM loads (set sizes 2, 3, 4, 5, and 6 stimuli). The set sizes were counterbalanced to ensure an even distribution of high and low set sizes, and no set size was repeated in consecutive trials. Images were randomly drawn from a session-respective stimulus pool, resulting in unique stimulus-response mapping for each participant.

### GNG task

We used openly available code from Zorowitz et al. (2025), but we modified the task in the following ways: 1) The timing of trials was modified so that the trial duration was fixed at 2.5s, and faster responses did not lead to an earlier onset of the next trial; 2) Attention checks were embedded evenly throughout the task (the original task did not include any). During attention checks, participants were required to press a key on a keyboard within 10 seconds; and 3) Only the conditions where points can be won (but not lost) were retained, that is, in each trial participants could win 1 or 10 points based on the choice. This modification was done based on our pilot studies and simulations and allowed us to decrease the total administration time while increasing the number of trials in the retained conditions, which improved parameter recovery.

Participants decided whether a robot appearing on a conveyor belt needed to be repaired: either repair the robot (press the spacebar) or do not repair it (withhold a response). The objective of the task was to correctly repair as many robots as possible. Participants learned which robots needed repair through trial and error, using probabilistic feedback: correct responses were rewarded probabilistically: 80% of correct and 20% of incorrect responses earned the participants 10 points, while 80% of incorrect and 20% of correct responses earned the participants only 1 point. The total points collected were converted into a performance bonus at the end of the task.

Participants performed 2 blocks with 80 trials each, resulting in 160 trials in total. The trial structure was pseudo-randomized and balanced such that there were 40 “go” robot types and 40 “no-go” robot types per block. There were 8 unique robot stimuli per block identified by a distinct chest plate symbol (stimulus). Each robot was repeated 8, 10 or 12 times (on average 10 times). There was a different stimulus set for each test session to prevent interference.

### Effort task

Participants performed 75 trials in total, sampled from a grid with 5 levels of reward and 5 levels of effort. Each combination of effort and reward was presented three times. The order of the trials was randomized for each participant. At the beginning of each trial, participants chose between rest and effort. If participants chose the rest option, they waited until their character walked to the coin and collected it. If they chose the effort option (level 1 to 5), they had a potential reward of (2 to 10 coins). However, participants had to run to the coin to reach it before the minotaur. Otherwise, no reward was granted. To run, participants were required to hold the right arrow and press the spacebar repeatedly until they reached the coin or the trial was over. The higher the effort level, the more spacebar presses were required to gain the coin. At the end of the experiment, the total number of collected coins was converted into a performance bonus.

To account for differences in physical ability between participants and ensure that the task difficulty was proportional to their physical ability, the task started with a titration procedure to calibrate effort levels based on how many button-presses each person could perform per a given unit of time. We measured each participant’s maximum number of button-presses within a fixed time window (6s; corresponding to the length of a trial). Specifically, they underwent a series of “run” trials against the minotaur, which became faster on every trial, until they failed to win twice in a row. The maximum number of presses that they made in the last winning trial was taken as their best performance. This measurement was then used to scale the effort required for each trial. This titration procedure was akin to what has been done in lab settings to measure physical effort by using maximal voluntary contraction (e.g., Lockwood et al., 2021).

### Foraging task

We modified the task from Harhen & Bornstein (2023) in the following ways: 1) In the original task, participants could occasionally (in 20% of the cases) switch to a planet with different properties after deciding to leave the current planet. In our task, the environment stayed stable until a break. Participants were informed that a switch happened at that point and nowhere else during the task; 2) We adjusted task parameters in the conditions, such as travel time to a next patch and reward depletion rate; 3) We embedded attention checks requiring participants to press a specific key on a keyboard within 10 seconds; 4) The original task had an “alien welcoming the participant” on every new planet. For the sake of time and to minimize distractions, we removed this element; 5) We changed the style of the instructions to align it with the instructions in the other tasks of the battery. Moreover, we edited slightly the wording of the comprehension quiz; and 6) Out of the original 4 conditions differing in the gem depletion rate and the travel time, we decided to keep only the 2 conditions with short and long travel times, while keeping the reward depletion rate constant. Our pilot studies revealed that the performance scores across conditions were highly correlated. Therefore, removing these conditions was not expected to influence the estimation of the exit threshold, which was our target variable in this task.

Participants performed two blocks of the task, each one with a fixed duration (6.5 minutes). The two blocks differed in travel time between planets (6s and 12s). The order of the blocks was randomized. Within each block, the travel time was constant and only changed when the next block started. Participants were informed that the total task time would be 13 minutes and that they were going to perform two runs with distinct travel times. However, they were not told the order of the runs.
